# TEV Induced Cleavage of Nuclei Tagged in Specific Cell Types (TIC-TaC) reveals histone profiles of activated neurons

**DOI:** 10.64898/2026.11.12.766233

**Authors:** Bradley J. Lukasak, Kai Shing, Callie E.W. Crawford, Jenna N. Beyer, Annabel K. Sangree, George Burslem, Erica Korb

## Abstract

Epigenomic profiling methods such as CUT&RUN/Tag are commonly employed to evaluate the genomic localization of histone modifications, transcription factors, and chromatin regulating proteins. Numerous transgenic mouse models also exist that label specific cell populations by leveraging Cre-based recombination and fluorescent reporters to allow for cell type specific epigenetic profiling *in vivo*. However, none of these systems allow for rapid, parallel processing for cell-type-specific CUT&RUN/Tag. Here, we report the development of **<u>T</u>**EV **I**nduced **<u>C</u>**leavage of Nuclei **<u>Ta</u>**gged in Specific **<u>C</u>**ell Types (TIC-TaC) in which we leverage a recombination-based system that results in conditional expression of a Sun1-GFP fusion protein with a TEV site introduced between Sun1 and GFP. Like the existing INTACT system, nuclei tagged by cell-type specific Cre recombinase activity can be efficiently purified using immunoprecipitation to allow for rapid, efficient, and in parallel nuclei isolation. However, with the addition of TEV protease, TIC-TaC nuclei can be subsequently removed from the beads allowing for downstream CUT&RUN/Tag. We applied this system to neurons isolated from the cortex to define the effects of seizure induced neuronal activation on numerous histone modifications and histone variant proteins and identified concordant shifts in H3K27ac, H3K27me3, H2A.Z and H2BE at seizure responsive genes. Together, through the development and application of TIC-TaC to allow for parallel processing of tissue for cell-type specific analysis, we uncover activity-dependent regulation of chromatin in the brain.

**Graphical Abstract:** 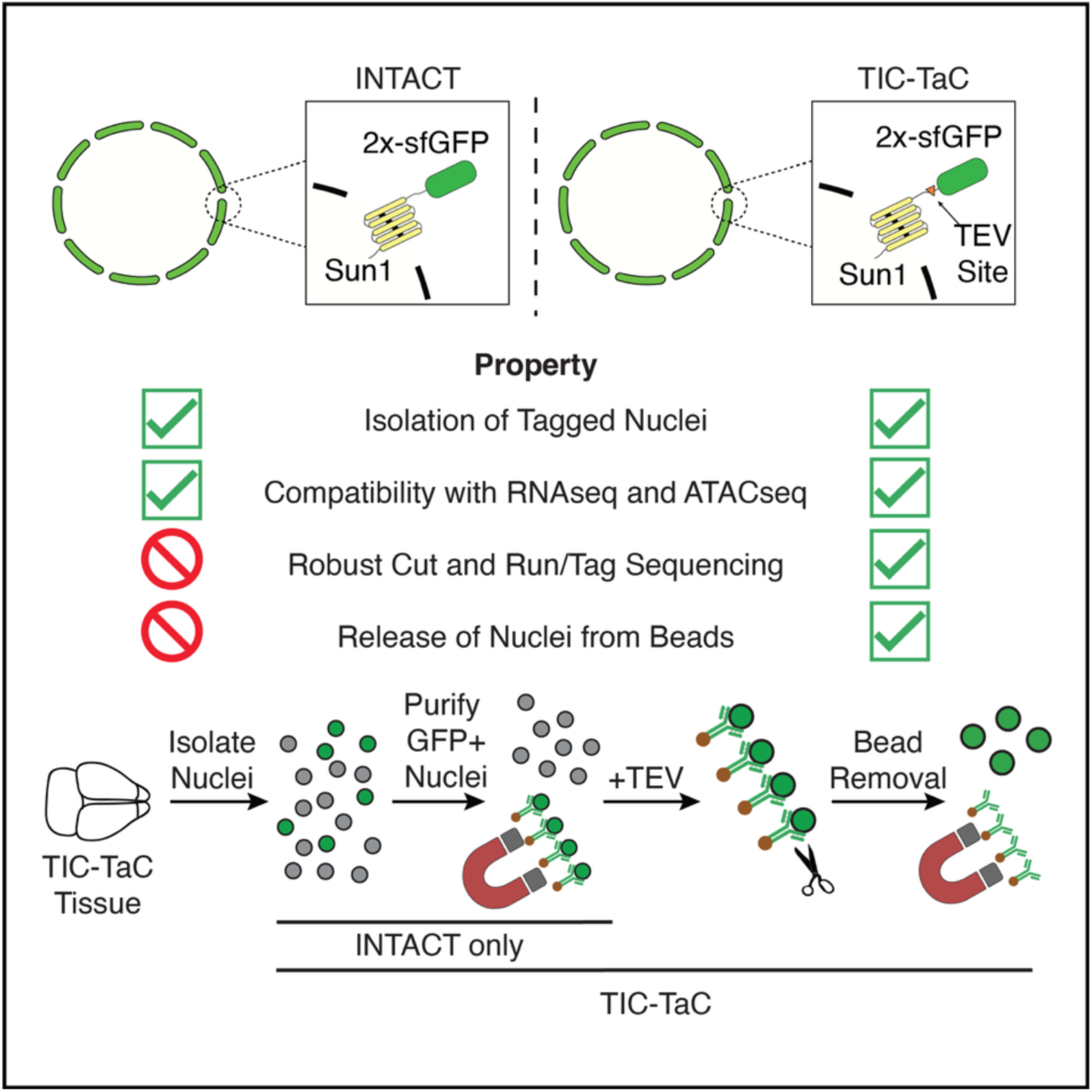

## Introduction

Chromatin, the physiologically relevant form of the eukaryotic genome, is a nucleoprotein complex consisting of DNA wrapped around two copies each of histones H2A, H2B, H3, and H4.^1^ Histone proteins are decorated with an extensive range of post-translation modifications (PTMs) that regulate chromatin structure and function.^2–5^ Such chromatin regulatory mechanisms are critical for establishing cell identity and allow for cellular function throughout the lifespan.^6,7^ Histone PTM signatures are also highly dynamic to allow for cellular responses to development signaling^8–11^, disease states^12–15^, environmental cues^16–18^, and more. However, rapid and efficient epigenetic profiling of multiple histone targets in a cell-type specific manner has proven challenging, limiting our understanding of dynamic epigenetic responses in complex tissue types.

The advent of CUT&RUN^19^ and subsequently CUT&Tag^20^ have allowed for epigenetic profiling with greater efficiency, at lower costs, and using lower cell numbers. Given the low input required, these approaches have the potential to allow for cell-type specific epigenetic profiling across many targets even from complex tissues, provided that specific cell types of interest can be readily isolated for downstream analysis. However, while powerful, cell sorting approaches are problematic as they introduce cell stress, are time consuming to apply across numerous conditions and replicates, can change gene expression,^21^ and can cause DNA damage.^22^ While single-cell CUT&Tag approaches have been developed and helped to define the chromatin states of individual cells^23^, they remain costly and thus out of reach to many, especially to apply across multiple replicates, conditions, and histone targets performed in parallel with sufficient sequencing depth. Thus, additional methods are required to make full use of the potential for CUT&RUN/Tag.

Isolation of nuclei tagged in specific cell types (INTACT) was developed as an alternative to cell sorting and single-cell approaches while still providing cell-type specificty.^24^ INTACT utilizes a conditionally expressed Sun1-GFP fusion protein that is expressed when paired with a specific Cre line to allow for cell type specific recombination. Upon recombination, Sun1-GFP is targeted to the nuclear lamina allowing for immunoprecipitation of the tagged nuclei using a GFP antibody and protein A (pA) fused magnetic beads. This method allows for in parallel processing of many samples at once, in a relatively short amount of time, with high efficiency and purity, and has been effectively paired with RNA-sequencing, ATAC-sequencing, ChIP-sequencing, and DNA methylation sequencing.^24–27^ Despite these advantages, INTACT is not fully compatible with downstream workflows for multiomic profiling. Because INTACT utilizes pA magnetic beads, any antibodies used in downstream assays will bind to the beads, reducing the effectiveness of approaches such as CUT&Tag and CUT&RUN. In addition, any assay that requires free nuclei that are not bound to beads cannot be performed following INTACT.

To address these challenges, we modified the INTACT system to allow for removal of the nuclei from beads under mild conditions, developing a method called **<u>T</u>**EV **I**nduced **<u>C</u>**leavage of Nuclei **<u>Ta</u>**gged in Specific **<u>C</u>**ell Types (TIC-TaC). In brief, we introduce a site for TEV protease between Sun1 and GFP to enable the cleavage of the fusion protein, separating nuclei from the beads. We validated that this system greatly improves CUT&RUN/Tag in nuclei isolated from tissue by applying it to neurons isolated from mouse brain. We then used TIC-TaC to define the seizure response of multiple histone modifications and histone variants to provide a map of the chromatin response to neuronal stimuli in the brain. We find that seizure-induced neuronal activity drives shifts in H3K27ac, H3K27me3, H2A.Z and H2BE that correspond to gene expression changes, demonstrating that these modifications and variants are highly dynamic in response to increased activity in cortical neurons. Together, these results establish TIC-TaC as an efficient tool for in parallel, cell-type-specific, chromatin profiling from highly complex biological tissue.

### Design

Given the power of the INTACT approach for providing efficient, inexpensive, and highly pure isolation of SUN1-GFP labelled nuclei, we sought to apply it for use in CUT&Tag applications. We confirmed that we were able to successfully perform INTACT and achieve highly pure GFP positive nuclei isolations from both cultured cells (**Figure S1A, S1B**) and from cells isolated from tissue in prior work.^28^ However, we found that using the resulting bead-bound nuclei for downstream CUT&Tag consistently failed. We speculated because nuclei are immobilized on protein A fused Dynabeads (pA-Dynabeads), downstream approaches that require antibodies will prove challenging. Antibodies will likely bind to pA-Dynabeads rather than their intended target given that these beads are designed for highly efficient antibody binding (**Figure S1C**). With no available methods to elute nuclei off the magnetic beads, we hypothesized that applications like CUT&Tag and CUT&RUN, that preserve nuclear structure and chromatin associations become ineffective as pA-Tn5 and MNase (used in CUT&Tag and CUT&RUN respectively) will be absorbed by bead-bound antibodies rather than reaching their intended chromatin targets (**Figure S1C**). To test this, we isolated Sun1-GFP transfected HEK293s using a standard INTACT method to isolate expressing nuclei through binding to pA-Dynabeads. Following isolation, the addition of an antibody targeting H3K27ac and a fluorescently labelled secondary antibody demonstrated the saturation of the Dynabeads with the antibodies (**Figure S1D**).

We therefore tried alternative methods to isolating GFP-positive nuclei. We first tested whether blocking additional bead binding sites improved downstream applications. We found that the addition of an IgG prior to antibody binding can effectively saturate pA binding sites (**Figure S1D**). However, this workflow proved incompatible with CUT&Tag (**Figure S1E**) likely because blocking pA-Dynabeads with an IgG provides pA-Tn5 with alternative binding partners (**Figure S1C**). Based on a recent study that used GFP-nanobody magnetic beads to perform INTACT^29^, we next evaluated whether nanobodies could provide a viable alternative for isolation of tagged nuclei and subsequent epigenomic profiling. We performed INTACT using nuclei isolated from the cortical tissue of mice expressing Sun1-GFP in excitatory neurons based on expression of Nex-Cre^30^. We observed that GFP-nanobody beads resulted in minimal enrichment of GFP positive nuclei (**Figure S1F-H**) and we also noted significant clumping of nuclei (**Figure S1I**), likely resulting in the inclusion of GFP negative nuclei. We note that fluorescence activated cell sorting (FACS) methods remain fully viable as an alternative to INTACT methods, but do not allow for the rapid and in-parallel isolation of many samples and conditions that we sought to achieve.

To address these limitations, we developed a new method that we termed **<u>T</u>**EV **I**nduced **<u>C</u>**leavage of Nuclei **<u>Ta</u>**gged in Specific **<u>C</u>**ell Types or ‘TIC-TaC’ (**Figure 1A-B**) to enable the elution of purified nuclei under gentle conditions. Within the linker between Sun1 and GFP, we introduced a TEV site that allows for proteolytic cleavage of the Sun1-GFP fusion and release of nuclei from magnetic beads. To minimize the impact of secondary structures in the fusion protein that could impact the proteolytic activity of TEV, we tested TEV sites in the N-terminal (TEVL) or C-terminal (LTEV) portion of the linker region that fuses Sun1 and GFP (**Figure S1J, S1K**). Using transfected HETK293T cells, nuclei were isolated using INTACT followed by the addition of TEV to release nuclei from beads. After three hours at 4C, both constructs effectively released nuclei from beads with LTEV performing slightly better (**Figure S1L-N**). As expected, in the absence of TEV, no nuclei were released to the supernatant.

**Figure 1.**
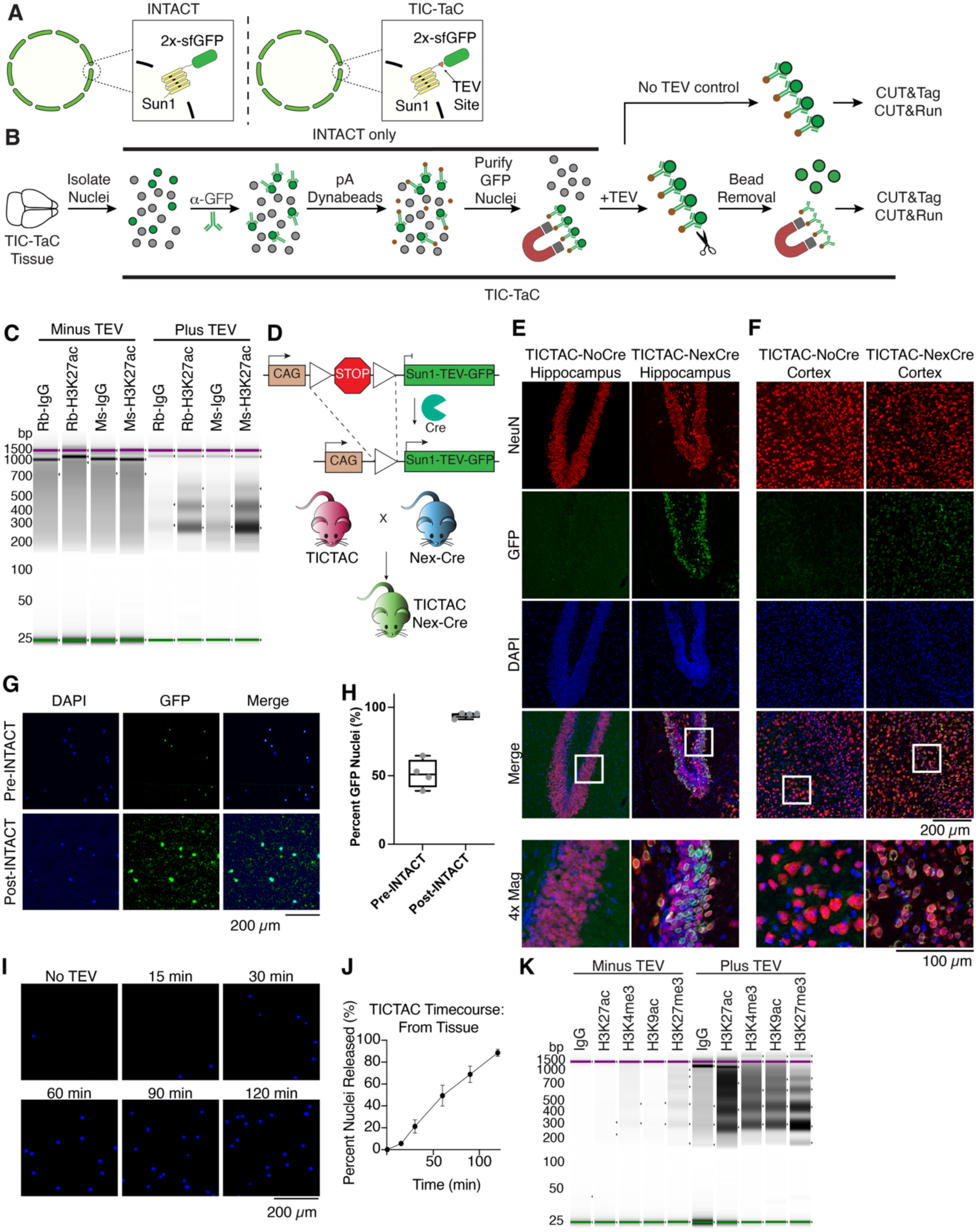
TIC-TaC principle and validation. **A)** Comparison of INTACT and TIC-TaC, highlighting TEV site in the linker region that enables release from beads under mild conditions. **B)** Schematic of experimental workflow for TIC-TaC. Following nuclei isolation, GFP positive nuclei are immobilized on pA-Dynabeads. Subsequent treatment with TEV removes nuclei from beads for downstream workflows. As a negative control, the same workflow is performed but without the addition of TEV prior to use for CUT&Tag/RUN. **C)** TapeStation of CUT&Tag from transfected HEK293 cells. On the left, samples remain on pA-Dynabeads for downstream CUT&Tag. On the right, samples were treated with TEV and then immobilized on ConA beads prior to CUT&Tag. IgG was used as a negative control. **D)** TIC-TaC mouse design, embedded at the Rosa26 locus. A CAG promoter drives conditional expression of Sun1-TEV-GFP following Cre-dependent excision of a floxed stop cassette. TIC-TaC mice were crossed to Nex-Cre mice for expression of Sun1-TEV-GFP in excitatory forebrain neurons. **E)** Staining for NeuN (red), GFP (green), and DAPI (blue) in the hippocampus of TIC-TaC^+/-^; Nex-Cre^-/-^ and TIC-TaC^+/-^; Nex-Cre^+/-^demonstrating Cre-dependent expression of Sun1-TEV-GFP of 7–8-week-old male mice. **F)** Staining for NeuN (red), GFP (green), and DAPI (blue) in the cortex of TIC-TaC^+/-^; Nex-Cre^-/-^and TIC-TaC^+/-^; Nex-Cre^+/-^demonstrating Cre-dependent expression of Sun1-TEV-GFP of 7–8-week-old male mice. **G)** Fluorescence imaging of pre-INTACT and post-INTACT purified cortical nuclei from TIC-TaC^+/-^; Nex-Cre^+/-^mice. **H)** Quantification of pre-INTACT and post-INTACT purified cortical nuclei from TIC-TIC-TaC^+/-^; Nex-Cre^+/-^mice demonstrating enrichment for GPF positive nuclei. **I)** Fluorescence imaging of nuclei following TEV-mediated elution off of pA-Dynabeads. Time course was conducted on nuclei from TIC-TaC^+/-^; Nex-Cre^+/-^mice over the course of two hours at RT. **J)** Quantification of nuclei following TEV-mediated elution off of pA-Dynabeads. Time course was conducted on nuclei from TIC-TaC^+/-^; Nex-Cre^+/-^mice over the course of two hours at RT. Demonstrating near quantitative recovery following a two-hour treatment. **K)** TapeStation of CUT&Tag in cortical nuclei purified from TIC-TaC^+/-^; Nex-Cre^+/-^mice. Minus TEV samples were left on pA-Dynabeads for downstream workflows and Plus TEV samples were bound to ConA beads following TEV-mediated elution. IgG was used as a negative control.

## Results

### TIC-TaC Allows for Recovery of Purified Nuclei for Downstream Applications

As a proof of principal, we proceeded to perform CUT&Tag from LTEV eluted nuclei (referred to as TIC-TaC from this point forward). Following TEV cleavage, eluted nuclei were immobilized on Concavalin A (ConA) beads for H3K27ac CUT&Tag, using both Rabbit and Mouse derived antibodies, selected due to well established roles in epigenetic regulatory pathways.^31,32^ As a control, we used the same nuclei and workflow but did not add TEV and used the resulting Dynabead-bound nuclei for CUT&Tag. We saw a clear increase in tagging efficiency for nuclei that were cleaved from beads using TEV as visualized on a TapeStation, regardless of the host the antibody was raised in (**Figure 1C**).

Following successful validation in cells, we generated a Cre-dependent TIC-TaC mouse by targeting the Rosa26 locus using homologous recombination (**Figure 1D**). Similarly to the INTACT model, the expression of Sun1-TEV-GFP is Cre-dependent by virtue of a floxed stop cassette. We validated our model by crossing the newly generated TIC-TaC line to Nex-Cre mice, producing TIC-TaC^+/-^; Nex-Cre^+/-^animals to achieve specific expression of Sun1-TEV-GFP in excitatory forebrain neurons.^33^ Immunohistochemistry (IHC) demonstrated a clear and robust Cre-dependent expression of Sun1-GFP in both the hippocampus and the cortex (**Figure 1E, 1F**). We further validated the model by performing a nuclei isolation from cortical tissue and purifying GFP positive nuclei (**Figure 1G, 1H**). One hemisphere of cortical tissue typically yields between 3 million and 6 million nuclei, of which approximately half are GFP positive (**Supplemental Table 1**).

The purified population was then eluted from the beads by the addition of TEV, with near complete release achieved after 2 hours at room temperature (**Figure 1I, 1J**). We note that elution of GFP positive nuclei derived from tissue takes longer than those derived from cells potentially due to more robust expression *in vivo* of the Sun1-GFP fusion following many weeks or months of accumulated expression.

Finally, we tested whether TIC-TaC allows for robust tagmentation in downstream CUT&Tag. Following immunoprecipitation of GFP positive nuclei, half received TEV protease to elute nuclei off the pA beads, while the other half remained bead bound as a control. Nuclei eluted from pA beads were then immobilized on Concanavalin A beads and CUT&Tag was performed for four histone modifications H3K27ac, H3K9ac, H3K27me3, and H3K4me3 (**Figure 1B**). We selected these marks due their well-established roles in gene regulation and to highlight the broad applicability of TIC-TaC.^31,34–39^ For every histone modification tested, TEV release of nuclei from the pA beads strongly increased signal (**Figure 1K**), demonstrating that removal of nuclei from pA beads substantially improves chromatin profiling. Together, these data indicate that Sun1-TEV-GFP mice allow for cell-type specific expression and subsequent elution from beads of targeted nuclei for downstream CUT&Tag.

### TIC-TaC achieves *in vivo* cell-type-specific histone modification profiling by CUT&Tag

Having demonstrated that TIC-TaC allows for robust nuclei elution and successful CUT&Tag library generation, we assessed its effectiveness via next generation sequencing. Marks associated with active transcription (H3K27ac, H3K9ac, H3K4me3), were enriched at neuronally expressed genes, whereas the signal for the repressive mark H3K27me3 was distributed across the gene body and at the TSS of lowly- or non-expressed neuronal genes as expected (**Figure 2A-D, S2A**). This was apparent at representative neuronal genes (*Grin2a, Syt1)* for H3K27ac, H3K9ac, and H3K4me3 (**Figure 2E, Figure S2A**). H3K27me3 was enriched across the gene bodies of genes not expected to be expressed in neurons like the oligodendrocyte transcription factor *Olig2,* the microglia enriched gene *Fcrls,* and the astrocyte enriched gene *Aldh1l1* (**Figure 2E**, **S2A**). Across loci, IgG signal was low, confirming Tn5 specificity (**Figure 2E, Figure S2A**). Together, these results demonstrate that TIC-TaC coupled with CUT&Tag enables high resolution profiling of histone modifications in defined cellular populations from complex biological tissue.

**Figure 2.**
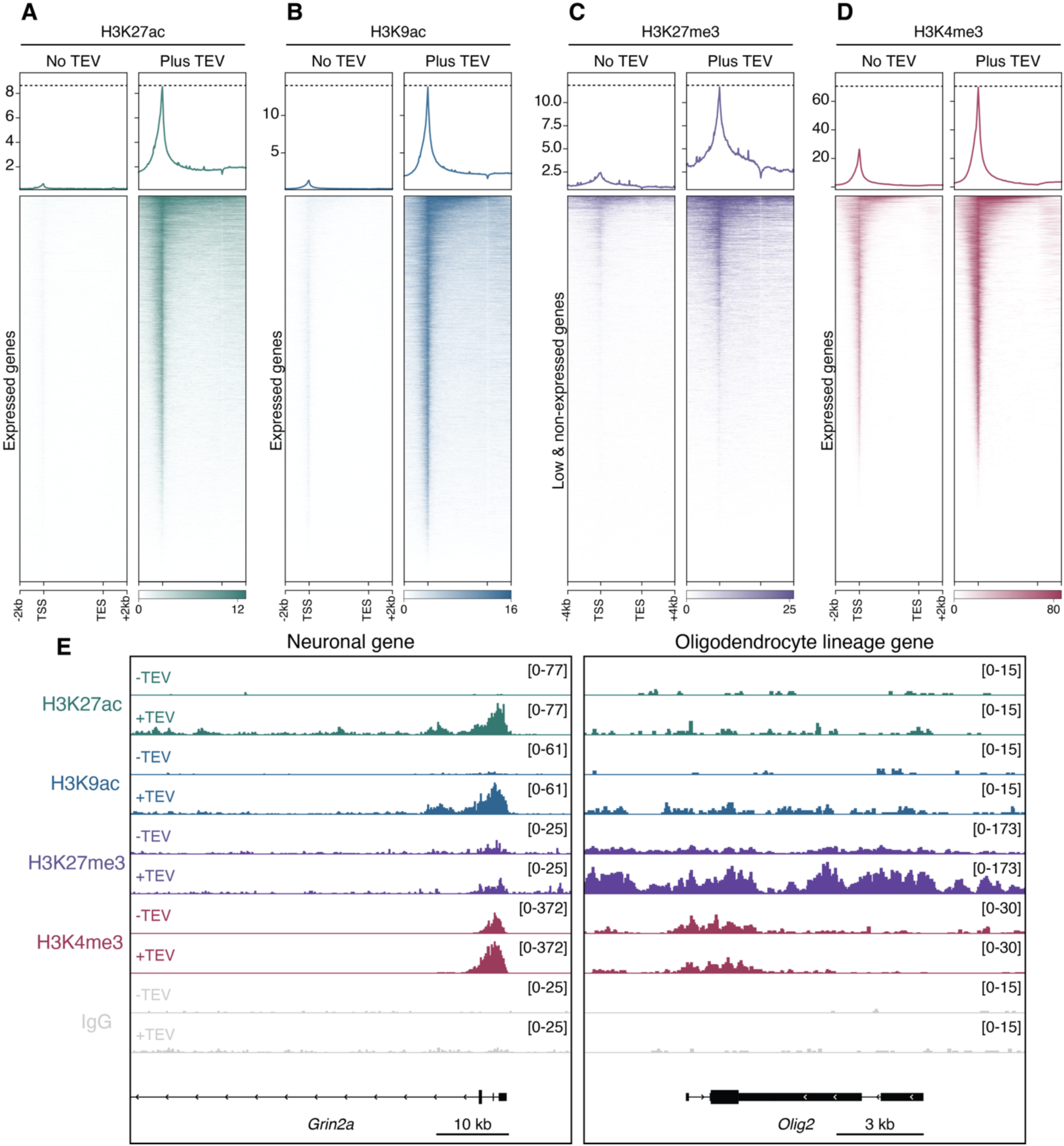
TIC-TaC achieves cell-type-specific histone modification profiling by CUT&Tag. **A)** Heatmap and metaplot show H3K27ac CUT&Tag signal at neuronally expressed genes for nuclei with no TEV treatment (left) and with TEV treatment (right) (n=16,266 genes). 2 kb upstream of the TSS and downstream of the TES are plotted. Data are merged across n=3 biological replicates. **B)** Heatmap and metaplot show H3K9ac CUT&Tag signal at neuronally expressed genes for nuclei with no TEV treatment (left) and with TEV treatment (right) (n=16,266 genes). 2 kb upstream of the TSS and downstream of the TES are plotted. Data are merged across n=3 biological replicates. **C)** Heatmap and metaplot show H3K27me3 CUT&Tag signal at low/non expressed neuronal genes for nuclei with no TEV treatment (left) and with TEV treatment (right) (n=6,649 genes). 4 kb upstream of the TSS and downstream of the TES are plotted. Data are merged across n=3 biological replicates. **D)** Heatmap and metaplot show H3K4me3 CUT&Tag signal at neuronally expressed genes for nuclei with no TEV treatment (left) and with TEV treatment (right) (n=16,266 genes). 2 kb upstream of the TSS and downstream of the TES are plotted. Data are merged across n=3 biological replicates. **E)** Representative CUT&Tag gene tracks for a neuronal marker gene (*Grin2a*, left) and non-neuronal marker gene (oligodendrocyte lineage, *Olig2*, right). Tracks show n=3 merged biological replicates.

### TIC-TaC achieves *in vivo* cell-type-specific histone modification profiling by CUT&RUN

Next, to determine whether TIC-TaC isolated nuclei are compatible with CUT&RUN, we applied the same workflow to profile H3K27ac, H3K9ac, H3K27me3, and H3K4me3, comparing nuclei with or without TEV mediated elution. Consistent with our CUT&Tag results, TEV mediated elution from pA beads resulted in globally enhanced signal across all marks tested using CUT&RUN (**Figure 3A-D, Figure S2B**). Notably, while H3K9ac and H3K4me3 displayed modest signal in the absence of elution, TEV protease treatment resulted in improved coverage for both marks (**Figure 3B, D, Figure S2B**). As with CUT&Tag, modifications associated with active transcription were enriched at the TSS of neuronally expressed genes in TEV eluted nuclei, whereas signal was markedly reduced in samples that remained bound to pA beads (**Figure 3A, B, D**). The repressive mark H3K27me3 displayed broad enrichment across the gene body and at the TSS of lowly or non-expressed neuronal genes in nuclei that were eluted by TEV (**Figure 3C**). These trends were recapitulated at single-locus resolution, where genome browser tracks at representative neuronal and non-neuronal loci (*Grin2a*, *Syt1, Olig2, Fcrl2s,* and *Aldh1l1*) again revealed TEV-dependent enrichment of expected marks and low IgG signal (**Figure 3E, Figure S2B**). Collectively, these results demonstrate that TEV mediated elution of isolated nuclei results in more efficient and consistent chromatin profiling by CUT&RUN. Further, we establish that TIC-TaC is compatible with multiple orthogonal chromatin mapping approaches for cell-type-specific histone modification profiling from tissue.

**Figure 3.**
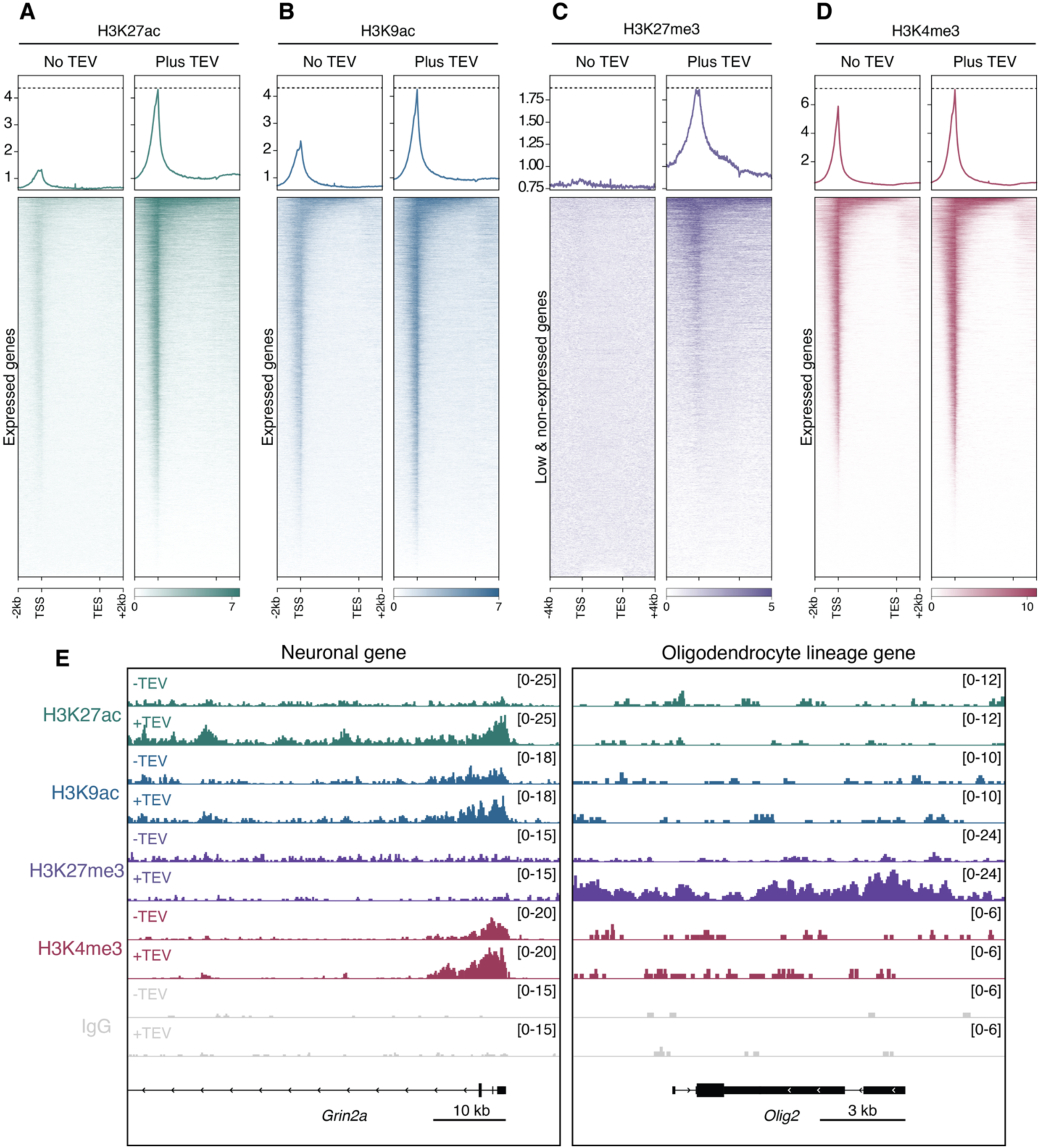
TIC-TaC achieves cell-type-specific histone modification profiling by CUT&RUN. **A)** Heatmap and metaplot show H3K27ac CUT&RUN signal at neuronally expressed genes for nuclei with no TEV treatment (left) and with TEV treatment (right) (n=16,266 genes). 2 kb upstream of the TSS and downstream of the TES are plotted. Data are merged across n=3 biological replicates. **B)** Heatmap and metaplot show H3K9ac CUT&RUN signal at neuronally expressed genes for nuclei with no TEV treatment (left) and with TEV treatment (right) (n=16,266 genes). 2 kb upstream of the TSS and downstream of the TES are plotted. Data are merged across n=3 biological replicates. **C)** Heatmap and metaplot show H3K27me3 CUT&RUN signal at low/non expressed neuronal genes for nuclei with no TEV treatment (left) and with TEV treatment (right) (n=6,649 genes). 4 kb upstream of the TSS and downstream of the TES are plotted. Data are merged across n=3 biological replicates. **D)** Heatmap and metaplot show H3K4me3 CUT&RUN signal at neuronally expressed genes for nuclei with no TEV treatment (left) and with TEV treatment (right) (n=16,266 genes). 2 kb upstream of the TSS and downstream of the TES are plotted. Data are merged across n=3 biological replicates. **E)** Representative CUT&RUN gene tracks for a neuronal marker gene (*Grin2a*, left) and non-neuronal marker gene (oligodendrocyte lineage, *Olig2*, right). Tracks show n=3 merged biological replicates.

### TIC-TaC chromatin profiling correlates with established bulk and cell-type-specific chromatin profiling methods

To benchmark our TIC-TaC CUT&Tag and TIC-TaC CUT&RUN data against existing chromatin profiling approaches, we compared it against three publicly available datasets which profiled H3K27ac, H3K27me3, and H3K4me3 using bulk CUT&RUN from 2-month-old mouse prefrontal cortex (**GSE222072**),^40^ CUT&RUN from FACS-isolated excitatory neuronal nuclei (**GSE218312**),^41^ and chromatin immunoprecipitation sequencing (ChIP-seq) from FACS-isolated neuronal nuclei (**GSE246777**).^42^ These datasets were chosen as they included both bulk and FACS-isolated nuclei preparations across multiple chromatin profiling methods.

Heatmaps of H3K27ac signal at neuronally expressed genes showed strong TSS enrichment across all five datasets as expected (**Figure 4A**). At the TSS, TIC-TaC CUT&Tag and TIC-TaC CUT&RUN correlated strongly with each other (**Figure S3A**). TIC-TaC CUT&Tag showed strong concordance with bulk cortex CUT&RUN, FACS CUT&RUN, and FACS ChIP-seq (**Figure 4B**). TIC-TaC CUT&RUN showed a similar pattern, correlating with bulk cortex CUT&RUN, FACS CUT&RUN, and FACS ChIP-seq (**Figure S3B**). Consistent with these observations, peak-level analysis identified 545 TSS peaks shared across all datasets (**Figure 4C**). These findings were further illustrated at *Grm5*, an excitatory neuronal gene encoding a metabotropic glutamate receptor, which showed similar TSS signal across all datasets (**Figure 4D**).

**Figure 4.**
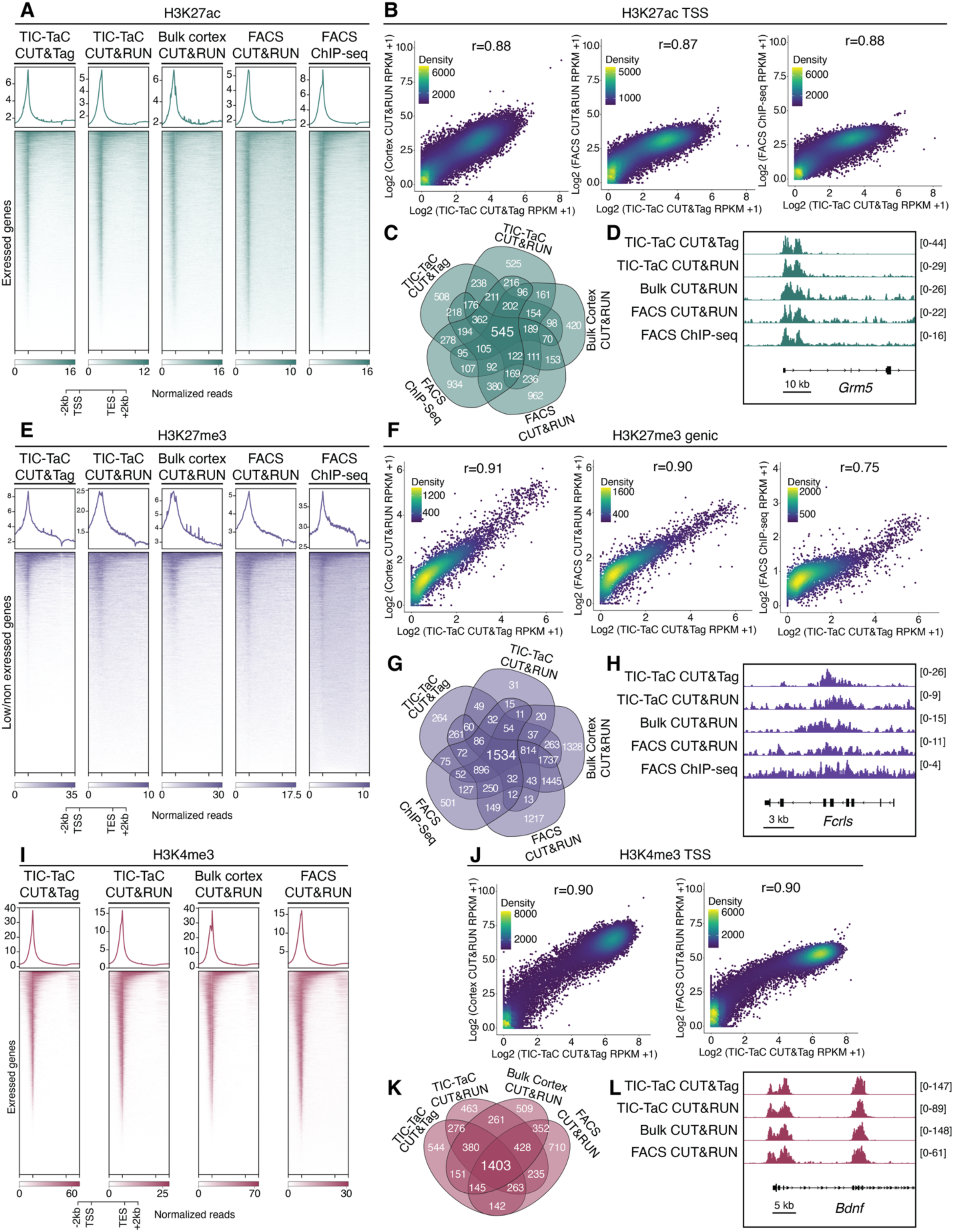
TIC-TaC chromatin profiling correlates with established bulk and cell-type-specific chromatin profiling methods. **A)** Heatmaps and metaplots show H3K27ac signal at neuronally expressed genes (n=16,266 genes). TIC-TAC CUT&Tag (n=3 replicates), TIC-TAC CUT&RUN (n=3 replicates), bulk cortex CUT&RUN (GSE222072, n= 2 replicates), FACS excitatory neuron CUT&RUN (GSE218312, n=2 replicates), FACS neuron ChIP-seq (GSE246772, n=1 replicate) are plotted. 2 kb upstream of the TSS and downstream of the TES are plotted. Data are merged across all biological replicates. **B)** Correlation between H3K27ac TSS reads from TIC-TAC CUT&Tag versus bulk cortex CUT&RUN (Spearman r=0.88, n=29,679), versus FACS excitatory nuclei CUT&RUN (Spearman r=0.87, n=29,679), and versus FACS neuronal nuclei ChIP-seq (Spearman r=0.88, n=29,679). **C)** Overlap of H3K27ac TSS peaks across TIC-TAC CUT&Tag, TIC-TAC CUT&RUN, bulk cortex CUT&RUN (GSE222072), FACS excitatory neuronal nuclei (GSE218312), and FACS neuronal nuclei ChIP-seq (GSE246772) datasets. **D)** Representative gene tracks showing H3K27ac TSS peaks at Grm5 across TIC-TAC CUT&Tag, TIC-TAC CUT&RUN, bulk cortex CUT&RUN (GSE222072), FACS excitatory neuronal nuclei (GSE218312), and FACS neuronal nuclei ChIP-seq (GSE246772) datasets. Data are merged across biological replicates. **E)** Heatmaps and metaplots show H3K27me3 signal at low/non expressed neuronal genes (n=6,649 genes). TIC-TAC CUT&Tag (n=3 replicates), TIC-TAC CUT&RUN (n=3 replicates), bulk cortex CUT&RUN (GSE222072, n= 3 replicates), FACS excitatory neuron CUT&RUN (GSE218312, n=2 replicates), FACS neuron ChIP-seq (GSE246772, n=1 replicate) are plotted. 2 kb upstream of the TSS and downstream of the TES are plotted. Data are merged across biological replicates. **F)** Correlation between H3K27me3 gene body reads from TIC-TAC CUT&Tag versus bulk cortex CUT&RUN (Spearman r=0.91, n=6,243), versus FACS excitatory nuclei CUT&RUN (Spearman r=0.90, n=6,243), and versus FACS neuronal nuclei ChIP-seq (Spearman r=0.75, n=6,243). **G)** Overlap of H3K27me3 gene body peaks across TIC-TAC CUT&Tag, TIC-TAC CUT&RUN, bulk cortex CUT&RUN (GSE222072), FACS excitatory neuronal nuclei (GSE218312), and FACS neuronal nuclei ChIP-seq (GSE246772) datasets. **H)** Representative gene tracks showing H3K27me3 gene body signal at Fcrls across TIC-TAC CUT&Tag, TIC-TAC CUT&RUN, bulk cortex CUT&RUN (GSE222072), FACS excitatory neuronal nuclei (GSE218312), and FACS neuronal nuclei ChIP-seq (GSE246772) datasets. Data are merged across biological replicates. **I)** Heatmaps and metaplots show H3K4me3 signal at neuronally expressed genes (n=16,266 genes). TIC-TAC CUT&Tag (n=3 replicates), TIC-TAC CUT&RUN (n=3 replicates), bulk cortex CUT&RUN (GSE222072, n= 2 replicates), and FACS excitatory neuron CUT&RUN (GSE218312, n=2 replicates. 2 kb upstream of the TSS and downstream of the TES are plotted. Data are merged across biological replicates. **J)** Correlation between H3K4me3 TSS reads from TIC-TAC CUT&Tag versus bulk cortex CUT&RUN (Spearman r=0.90, n=29,679) and versus FACS excitatory nuclei CUT&RUN (Spearman r=0.90, n=29,679). **K)** Overlap of H3K4me3 TSS peaks across TIC-TAC CUT&Tag, TIC-TAC CUT&RUN, bulk cortex CUT&RUN (GSE222072) and FACS excitatory neuronal nuclei (GSE218312) datasets. **L)** Representative gene tracks showing H3K4me3 TSS peaks at *Bdnf* across TIC-TAC CUT&Tag, TIC-TAC CUT&RUN, bulk cortex CUT&RUN (GSE222072), and FACS excitatory neuronal nuclei (GSE218312) datasets. Data are merged across biological replicates.

For the repressive mark H3K27me3, heatmaps show gene body enrichment with a peak at the TSS for lowly or non-expressed neuronal genes (**Figure 4E**). H3K27me3 at the gene body and TSS were strongly correlated between TIC-TaC CUT&Tag and TIC-TaC CUT&RUN (**Figure S3A**). At the gene body, TIC-TaC CUT&Tag correlated strongly with bulk cortex CUT&RUN, FACS CUT&RUN, and FACS ChIP-seq (**Figure 4F**), with TIC-TaC CUT&RUN showing a similar pattern (**Figure S3C**). At the TSS, correlations followed a comparable trend for both TIC-TaC CUT&Tag (**Figure S3D**) and TIC-TaC CUT&RUN (**Figure S3E**) against each dataset. Peak overlap analysis identified a large core set of 1,534 H3K27me3 domains shared across methods (**Figure 4G**). Genome browser tracks at the microglial enriched gene *Fcrls* showed robust H3K27me3 enrichment across the gene body for each dataset, consistent with its repression in neurons (**Figure 4H**).

Finally, comparison of H3K4me3 profiles at the TSS also revealed strong concordance between TIC-TaC CUT&Tag and TIC-TaC CUT&RUN datasets (**Figure S3A**). TIC-TaC CUT&Tag was highly correlated with bulk cortex CUT&RUN and FACS CUT&RUN at the TSS (**Figure 4J**). Similarly, TIC-TaC CUT&RUN correlated with bulk cortex CUT&RUN and FACS CUT&RUN at the TSS (**Figure S3F**). Peak calling and overlap analysis identified 1,403 TSS peaks shared across all datasets (**Figure 4K**). At the neuronal gene *Bdnf*, TIC-TaC CUT&Tag/RUN recapitulated the sharp TSS enrichment observed in bulk and FACS CUT&RUN datasets (**Figure 4L**). Together, these data demonstrate that TIC-TaC CUT&Tag and TIC-TaC CUT&RUN produce histone PTM chromatin profiles that are highly concordant with established bulk and cell-type-specific profiling methods, capturing both active (H3K27ac, H3K4me3) and repressive (H3K27me3) chromatin environments with high reproducibility. However, in addition to achieving similarly robust data as the methods used to generate these datasets, TIC-TaC provides cell-type specificity (unlike bulk tissue methods) while also allowing for in-parallel, rapid, and highly efficient nuclei purification (unlike FACS-based methods). Together, these data indicate that TIC-TaC provides robust sequencing data in line with prior approaches with the added benefits of efficient cell-type-specific profiling that can be performed in parallel across samples in place of cell sorting.

### TIC-TaC CUT&Tag captures activity dependent remodeling of histone modifications in excitatory neurons

Next, we sought to determine whether TIC-TaC CUT&Tag can resolve dynamic, cell-type-specific chromatin changes in response to a stimulus. We induced seizures in TIC-TaC^+/-^; Nex-Cre^+/-^mice via intraperitoneal injection of pentylenetetrazol (PTZ), which drives widespread increases in excitatory neuronal activity through inhibition of GABAA receptors.^43–45^ Nuclei were isolated from cortical tissue using TIC-TaC from home cage control mice and PTZ-treated mice and CUT&Tag was performed for H3K27ac, H3K9ac, H3K4me3, H3K27me3, and IgG (**Figure 5A**) again selected due their well-established roles in transcription as well as their importance in neuron identity, neuronal function, and behavior/memory.^46–60^ We observed the expected enrichment patterns in both home cage and seizure conditions with TSS enrichment for H3K27ac, H3K9ac, and H3K4me3 at neuronally expressed genes, and TSS and gene body enrichment for H3K27me3 at lowly or non-expressed neuronal genes (**Figure 5B-E**). Following seizure, H3K27ac was globally increased at the TSS of neuronally expressed genes (**Figure 5B, E**), while H3K9ac and H3K4me3 showed modest global changes (**Figure 5C**). The repressive mark H3K27me3 increased at the TSS and gene body of lowly or non-expressed neuronal genes (**Figure 5D)**. Confirming these results, we observed increased spike-in normalized signal at the TSS for H3K27ac, H3K9ac, H3K27me3, and H3K4me3 (**Figure 5F-I**). Together, these data establish that TIC-TaC coupled with CUT&Tag resolves activity dependent chromatin dynamics across multiple histone PTMs from a defined population of cortical nuclei.

**Figure 5.**
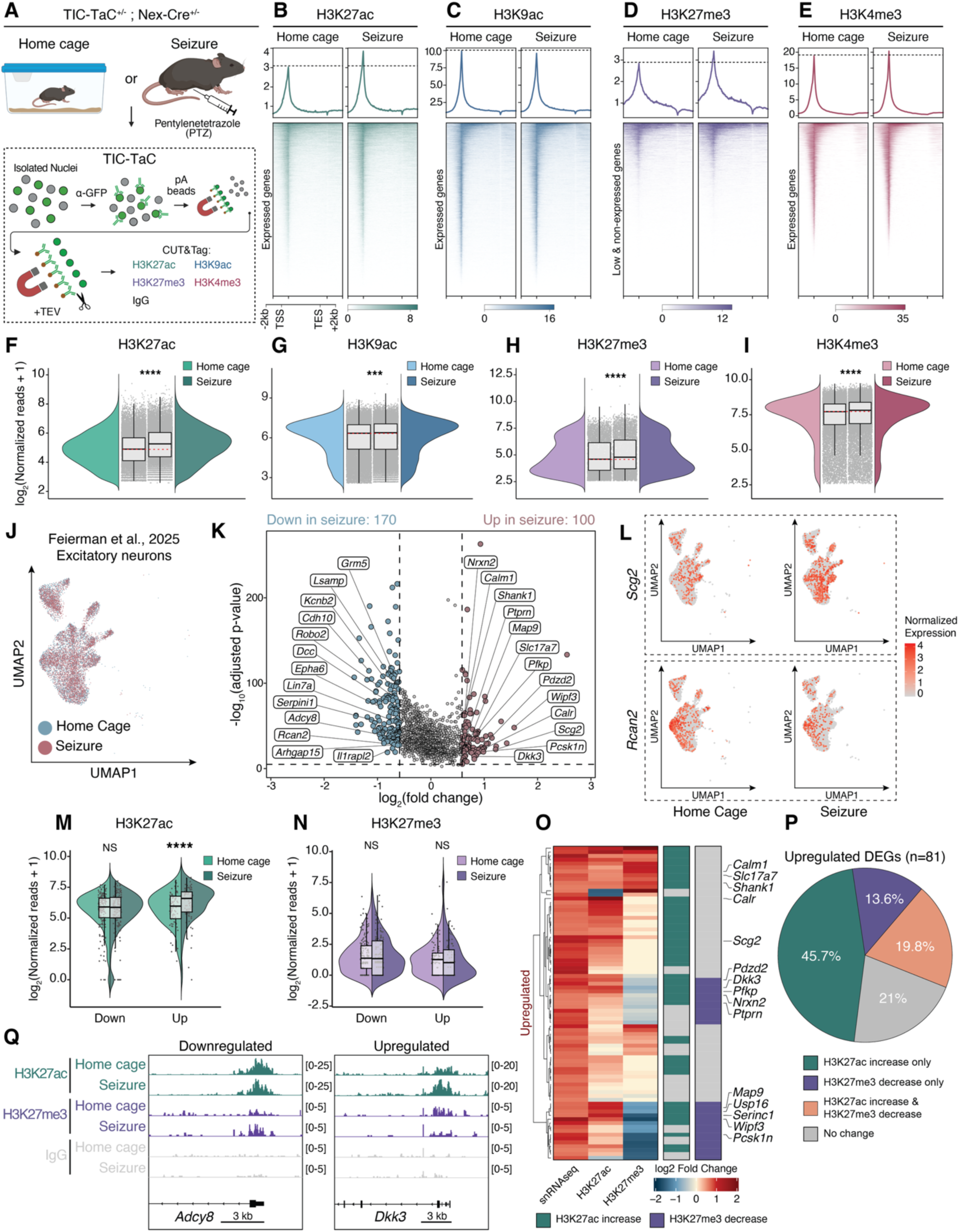
TIC-TaC CUT&Tag captures activity dependent remodeling of histone modifications in excitatory neurons. **A)** Schematic showing experimental overview. Cortical nuclei from home cage (n=2) or PTZ treated (n=3) mice were isolated by TIC-TaC. From each mouse, CUT&Tag was performed for H3K27ac, H3K9ac, H3K4me3, H3K27me3, and IgG in parallel. **B)** Heatmap and metaplot show H3K27ac CUT&Tag signal at neuronally expressed genes for nuclei isolated from home cage (left) and seizure (right) groups (n=16,266 genes). 2 kb upstream of the TSS and downstream of the TES are plotted. Data are merged across n=3 biological replicates for home cage mice and n=2 biological replicates for seizure-induced mice. **C)** Heatmap and metaplot show H3K9ac CUT&Tag signal at neuronally expressed genes for nuclei isolated from home cage (left) and seizure (right) groups (n=16,266 genes). 2 kb upstream of the TSS and downstream of the TES are plotted. Data are merged across n=3 biological replicates for home cage mice and n=2 biological replicates for seizure-induced mice. **D)** Heatmap and metaplot show H3K27me3 CUT&Tag signal at low/non neuronally expressed genes for nuclei isolated from home cage (left) and seizure (right) groups (n=6,649 genes). 2 kb upstream of the TSS and downstream of the TES are plotted. Data are merged across n=3 biological replicates for home cage mice and n=2 biological replicates for seizure-induced mice. **E)** Heatmap and metaplot show H3K4me3 CUT&Tag signal at neuronally expressed genes for nuclei isolated from home cage (left) and seizure (right) groups (n=16,266 genes). 2 kb upstream of the TSS and downstream of the TES are plotted. Data are merged across n=3 biological replicates for home cage mice and n=2 biological replicates for seizure-induced mice. **F)** Violin plots show CUT&Tag spike in normalized TSS H3K27ac reads. Statistics are two-sided Wilcoxon ranked-sum tests (n=15,812, ****p<0.0001). **G)** Violin plots show CUT&Tag spike in normalized TSS H3K9ac reads. Statistics are two-sided Wilcoxon ranked-sum tests (n=20,893, ***p<0.001). **H)** Violin plots show CUT&Tag spike in normalized TSS H3K27me3 reads. Statistics are two-sided Wilcoxon ranked-sum tests (n=6,258, ****p<0.0001). **I)** Violin plots show CUT&Tag spike in normalized TSS H3K4me3 reads. Statistics are two-sided Wilcoxon ranked-sum tests (n=16,674, ****p<0.0001). **J)** Uniform manifold approximation and projection (UMAP) for dimension reduction of single-nucleus transcriptomic profiles from adult male (2–4 months) mouse cortices (n=2 home cage, n=3 seizure). Excitatory neuron clusters are shown for home cage control (blue) and seizure (red) groups. **K)** Volcano plot shows excitatory neuron differentially expressed genes (FC>1.5, Bonferroni-corrected P<0.00001) identified by MAST in seizure compared to home cage control nuclei. **L)** Feature plots showing normalized expression in excitatory neurons for representative upregulated gene *Scg2* (top) and downregulated gene *Rcan2* (bottom). **M)** Violin plots show CUT&Tag normalized reads for H3K27ac binned by downregulated (left, n=158) and upregulated genes (right, n=81). Statistics are two-sided Wilcoxon ranked-sum tests; ****p<0.0001, ns = not significant. **N)** Violin plots show CUT&Tag normalized reads for H3K27me3 binned by downregulated (left, n=158) and upregulated genes (right, n=81). Statistics are two-sided Wilcoxon ranked-sum tests; ns = not significant. **O)** Heatmap shows excitatory neuron upregulated genes (n=81) in seizure compared to home cage control nuclei. Log2 fold changes for gene expression, H3K27ac, and H3K27me3 are shown. Bars to right of heatmap indicate direction of histone modification change (1.2-fold change, teal: H3K27ac increase, purple: H3K27me3 decrease). **P)** Pie chart shows percentages of upregulated genes that have increased TSS H3K27ac only (green), decreased TSS H3K27me3 only (purple), increased TSS H3K27ac and decreased TSS H3K27me3 (orange), or exhibit no change in H3K27ac or H3K27me3 (grey). **Q)** Gene tracks show representative downregulated (*Adcy8*, left) and upregulated (*Dkk3*, right) genes and accompanying changes in H3K27ac and H3K27me3.

To evaluate whether the chromatin changes identified by TIC-TaC into CUT&Tag correspond to transcriptional differences, we integrated our data with previously generated single-nucleus transcriptomic profiles from home-cage and seizure-induced mice.^45^ MAST differential gene expression analysis identified 270 seizure responsive genes, with 170 downregulated and 100 upregulated using a strict cutoff to limit false positives (1.5-fold change, p<0.00001) (**Figure 5J, K**). Feature plots confirmed increased expression of *Scg2*, which is involved in packaging neuropeptides into secretory vesicles,^61^ as well as decreased expression of *Rcan2* which inhibits calcineurin to regulate calcium signaling in neurons (**Figure 5L**).^62^ To independently validate MAST differential gene expression, we performed a more stringent pseudobulk analysis, which identified 44 downregulated and 33 upregulated genes in seizure compared to home-cage excitatory neurons (**Figure S4A**). Comparison of genes identified by MAST and pseudobulk revealed 27 shared downregulated genes (**Figure S4B**) and 17 shared upregulated genes (**Figure S4B**). Feature plots confirmed increased expression of *H3f3b*, which encodes the activity regulated histone variant H3.3,^63^ as well as decreased expression of *Htr1f* which encodes the seizure responsive 5-HT1F serotonin receptor (**Figure S4C**).^64^

We next examined whether these transcriptional changes were accompanied by corresponding changes in histone PTMs by comparing normalized CUT&Tag reads at the TSS of downregulated, upregulated, and unchanged genes. TSS H3K27ac signal was significantly increased with seizure at upregulated genes whereas downregulated genes showed no change (**Figure 5M**). H3K27me3 trended towards a decrease at upregulated genes and was unchanged at downregulated genes (**Figure 5N**). H3K9ac and H3K4me3 signal at the TSS were increased at both downregulated and upregulated genes (**Figure S4D-E**). These findings support a model in which multiple modifications synergize to affect gene expression with H3K27ac increasing in both stable and upregulated genes while H3K27me3 counteracts its effects to retrain gene expression changes at the set of non-differentially expressed genes.

Gene set enrichment analysis (GSEA) using CUT&Tag fold-changes as a ranking metric further supported these patterns. H3K27ac showed the strongest enrichment among all four marks (**Figure S4F**). Downregulated genes were significantly enriched for decreased H3K27ac, and upregulated genes were significantly enriched for increased H3K27ac (**Figure S4F**). H3K27me3 showed a negative enrichment at upregulated genes and a positive enrichment at downregulated genes although these did not reach statistical significance (**Figure S4F**). H3K9ac and H3K4me3 displayed moderate positive enrichment and were not statistically significant across upregulated or downregulated gene sets (**Figure S4F**). Together, these results suggest that H3K27ac and H3K27me3 dynamics most closely tracked with seizure induced transcriptional changes in excitatory neurons, consistent with their established roles in transcriptional activation and repression, respectively, while H3K9ac and H3K4me3 were largely uncoupled from seizure induced transcriptional changes.

Next, we compared the log2 fold change of these genes with the log2 fold change of H3K27ac and H3K27me3 (**Figure 5O**). Genes with concordant epigenetic and transcriptional changes included synaptic and activity-regulated genes such as *Calm1, Slc17a7*, *Shank1*, *Calr*, *Scg2*, *Nrxn2*. Strikingly, 45.7% of upregulated genes displayed a concordant change in H3K27ac signal (greater than 20% increase; **Figure 5P**), while a smaller percentage (13.6%) displayed a concordant change in H3K27me3 signal (greater than 20% *decrease*; **Figure 5P**). Further, 19.8% of upregulated genes displayed concordant changes in both H3K27ac and H3K27me3 at the TSS (**Figure 5P**). Consistent with these trends, *Dkk3,* which is transcriptionally upregulated, exhibits increased H3K27ac signal and decreased H3K27me3 signal at the TSS (**Figure 5Q**). In alignment with our data demonstrating that downregulated genes display more modest global changes in H3K27 modifications, only 14.6% of genes showed decreased H3K27ac and 27.2% of genes showed increased H3K27me3 signal at the TSS (**Figure S4G**). Downregulated genes displaying concordant epigenetic and transcriptional changes included calcium signaling and synaptic genes *Adcy8*, *Rcan2*, *Grm5*, *Epha6*, *Kcnb2*, and *Robo2* (**Figure 5Q, Figure S4G**).

A substantial fraction of non-differentially expressed genes showed significant changes in histone PTM signal despite unchanged transcript level (**Figure S4H**). The largest subsets showed increased H3K27ac only (21.5%), increases in both H3K27ac and H3K27me3 (19.8%), and increased H3K27ac and decreased H3K27me3 (16.2%) (**Figure S4H**). 11.3% of non-DEGs showed no changes in H3K27ac or H3K27me3 signal (**Figure S4H**). At representative downregulated (*Adcy8*, *Arhgap15*) and upregulated (*Dkk3*, *Wipf3*) loci, H3K27ac and H3K27me3 displayed concordant seizure associated changes, while H3K9ac and H3K4me3 were more limited (**Figure S4I**). Taken together, these results reveal concordant changes in H3K27ac and H3K27me3 at seizure-responsive genes in neurons and suggest a model in which multiple modifications act to both promote and restrain gene expression changes.

### TIC-TaC CUT&Tag reveals activity dependent remodeling of histone variants in excitatory neurons

Finally, we tested the effect of seizure induced neuronal activity on the histone variants H2A.Z and H2BE, which have been demonstrated to have important roles in modulating neuronal plasticity.^28,65^ We induced seizures to drive neuronal activation using the same paradigm described above, and isolated excitatory cortical neurons to perform TIC-TaC followed by CUT&Tag for H2A.Z and H2BE. We observed a global increase in H2A.Z and decrease in H2BE at neuronally expressed genes following seizure induced neuronal activity (**Figure 6A-B**). Quantification of CUT&Tag reads confirmed that H2A.Z was increased and H2BE was decreased at the TSS (**Figure 6C-D**). We next tested whether these changes corresponded seizure induced gene expression in excitatory neurons.^28^ Because H2BE typically promotes an open chromatin signal and H2A.Z can repress transcription, we considered changes to be concordant if H2BE increased and H2A.Z decreased at genes with increased expression and vice versa. H2A.Z signal was significantly increased at both downregulated and upregulated genes, while H2BE signal was significantly decreased at downregulated genes (**Figure 6E-F**). Downregulated genes that gained H2A.Z or lost H2BE included ion channels (*Kcnq3*, *Kcnip4*, *Nalcn*), neurotransmitter receptors (*Gabrb2*, *Gabrg3*, *Grid2*, *Grm3*, *Gria4*, *Htr1f*, Neto1) and synaptic adhesion and organization (*Cdh2*, *Nrxn1*, *Cadm2*, *Cntn1*, *Csmd1*) (**Figure 6G**). Upregulated genes that lost H2A.Z or gained H2BE included genes related to synaptic organization (*Nptxr*, *Nrxn2*, *Ptprn*, *Shank1*, *Slc17a7*), vesicle and neuropeptide secretion (*Chgb*, *Pcsk1n*, *Ptprn*, *Scg3*), metabolism (*Apoe*, *Cox6c*, *Ociad2*, *Osbpl3*, *Serinc1*, *Slc2a3*), the histone variant H3.3 (*H3f3b*), and the histone methyltransferase *Dot1l* (**Figure 6H**). Among downregulated genes, the largest fraction lost H2BE (29.1%), while 18.4% only gained H2A.Z and 13.9% exhibited both changes (**Figure 6I**). Similarly, among upregulated genes, 37% gained H2BE whereas smaller proportions only lost H2A.Z (8.6%) or exhibited both changes (2.5%) (**Figure 6J**). As with histone PTMs, a large portion of non-DEGs exhibited changes in histone variant signal upon seizure induce neuronal activation (**Figure S5A-B**). Representative gene tracks show the downregulated AMPAR subunit gene *Gria4* which gains H2A.Z and loses H2BE, and the upregulated protein tyrosine phosphatase receptor *Ptprn* which loses H2A.Z and gains H2BE (**Figure 6K**). Further, our results recapitulate previously reported stimulus induced changes in H2A.Z binding in the brain where H2A.Z is decreased at *Egr1* and increased at *Ppp3ca* (**Figure S5C**).^65^ Among downregulated genes, a large cluster lost H2BE and a subset of these genes further gained H2A.Z or H3K27me3 or lost H3K27ac.

**Figure 6.**
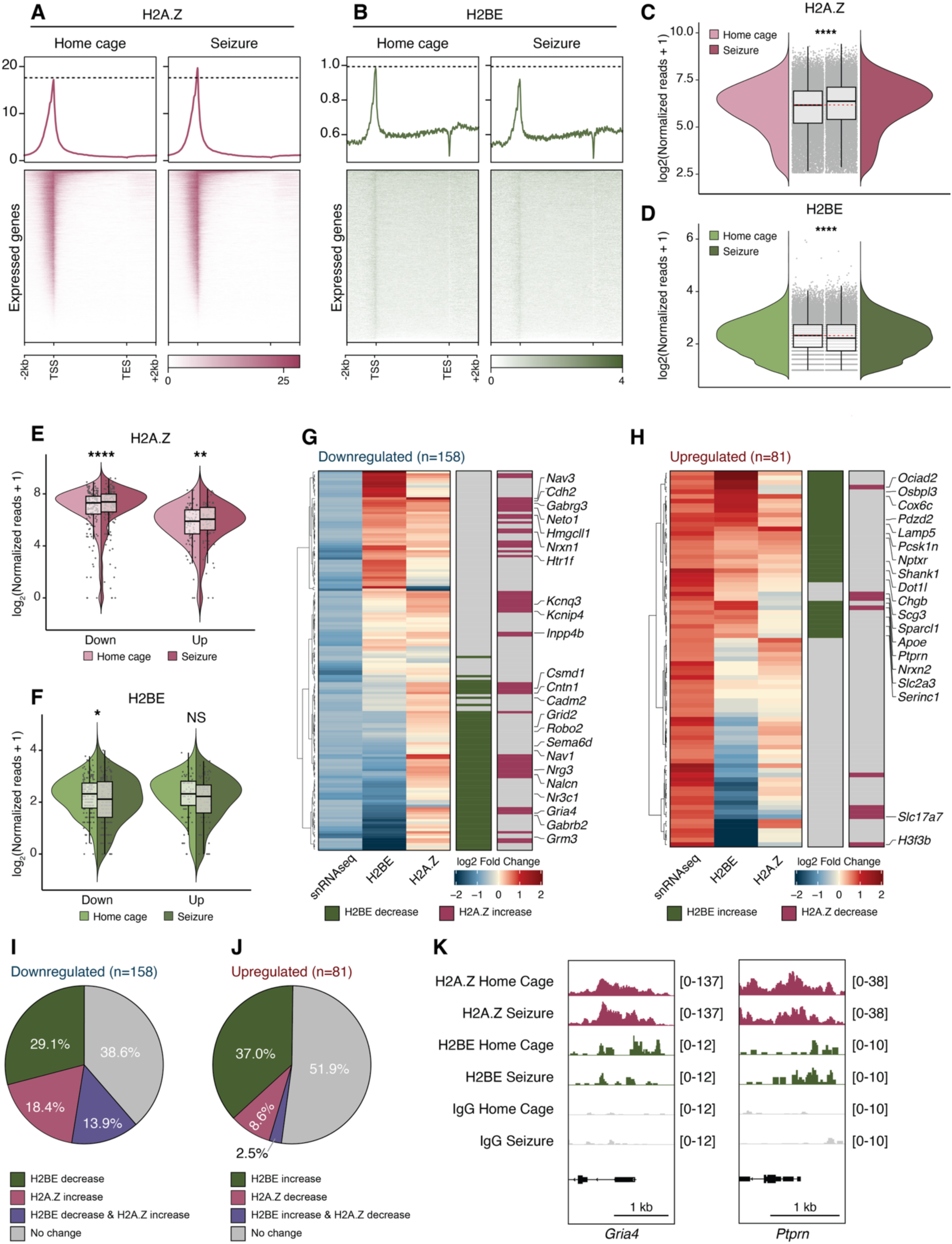
TIC-TaC CUT&Tag captures activity dependent remodeling of histone variants in excitatory neurons. **A)** Heatmap and metaplot show H2A.Z CUT&Tag signal at neuronally expressed genes for nuclei isolated from home cage (left) and seizure (right) groups (n=16,266 genes). 2 kb upstream of the TSS and downstream of the TES are plotted. Data are merged across n=3 biological replicates for home cage mice and n=3 biological replicates for seizure-induced mice. **B)** Heatmap and metaplot show H2BE CUT&Tag signal at neuronally expressed genes for nuclei isolated from home cage (left) and seizure (right) groups (n=16,266 genes). 2 kb upstream of the TSS and downstream of the TES are plotted. Data are merged across n=3 biological replicates for home cage mice and n=3 biological replicates for seizure-induced mice. **C)** Violin plots show CUT&Tag TSS H2A.Z reads. Statistics are two-sided Wilcoxon ranked-sum tests (n= 18,675, ****p<0.0001). **D)** Violin plots show CUT&Tag spike in normalized TSS H2BE reads. Statistics are two-sided Wilcoxon ranked-sum tests (n= 22,924, ****p<0.0001). **E)** Violin plots show CUT&Tag H2A.Z TSS reads binned by downregulated (left, n=158) and upregulated genes (right, n=81). Statistics are two-sided Wilcoxon ranked-sum tests; ****p<0.0001, **p<0.01. **F)** Violin plots show CUT&Tag TSS H2BE reads binned by downregulated (left, n=158) and upregulated genes (right, n=81). Statistics are two-sided Wilcoxon ranked-sum tests; *p<0.05, ns = not significant. **G)** Heatmap shows excitatory neuron downregulated genes (n=158) in seizure compared to home cage control nuclei. Log2 fold changes for gene expression, H2BE, and H2A.Z are shown. Bars to right of heatmap indicate direction of histone variant change (1.2-fold change, green: H2BE decrease, maroon: H2A.Z increase). **H)** Heatmap shows excitatory neuron upregulated genes (n=81) in seizure compared to home cage control nuclei. Log2 fold changes for gene expression, H2BE, and H2A.Z are shown. Bars to right of heatmap indicate direction of histone variant change (1.2-fold change, green: H2BE increase, maroon: H2A.Z decrease). **I)** Pie chart shows percentages of downregulated genes that have decreased TSS H2BE only (green), increased TSS H2A.Z only (maroon), increased TSS H2A.Z and decreased TSS H2BE (purple) or no change in H2A.Z or H2BE (grey). **J)** Pie chart shows percentages of upregulated genes that have increased TSS H2BE only (green), decreased TSS H2A.Z only (maroon), increased TSS H2BE and decreased TSS H2A.Z (purple), or no change in H2A.Z or H2BE (grey). **K)** Representative genome browser tracks show home cage and seizure conditions for H2A.Z and H2BE at downregulated (*Gria4,* left) and upregulated (*Ptprn*, right) genes.

However, the pattern of changes in H2A.Z, H3K27ac and H3K27me3 was largely independent of H2BE (**Figure S5D**). Among upregulated genes, increased H3K27ac was the dominant activity-induced change, with a subset of these genes also gaining H2BE or losing H3K27me3, and H2A.Z loss occurring less frequently (**Figure S5E**). Together, these data demonstrate that seizure induced neuronal activity remodels histone variant occupancy in excitatory neurons, driving global loss of H2BE and gain of H2A.Z, and locus specific changes that correlate with transcription. Further, this work demonstrates that TIC-TaC can successfully be used to efficiently profile chromatin across multiple replicates, conditions, modifications, and histone variants.

## Discussion

In this study, we developed TIC-TaC, a method that allows for reliable and reproducible epigenomic profiling of specific cell populations *in vivo*. Previously, epigenomic profiling of specific cell types required cell sorting approaches. While powerful, cell sorting is time consuming, stressful on cells, and renders in-parallel sample processing impractical. We demonstrated that the addition of a TEV site to the transgene previously used for INTACT purifications allows for nuclei elution for downstream CUT&RUN and CUT&Tag. This TIC-TaC approach allows for complete in-parallel processing of large sample pools in approximately 4 hours, reducing batch affects that can arise when processing individual samples sequentially.

We found that TIC-TaC greatly improved CUT&Tag and CUT&RUN signal compared to nuclei left on pA Dynabeads. However, in a few cases we were able to detect signal in the absence of TEV elution. Particularly for H3K4me3 and H3K27ac when performing CUT&RUN. This may be due to epitope abundance or antibody affinity allowing for some Tn5 targeting to overcome Dynabead interference for the very robust H3K4me3 and H3K27ac. While the binding affinity for pA Dynabeads is strong, it could be partially overcome by an antibody that has a comparable affinity for its epitope. Notably, even in these circumstances, TIC-TaC substantially improved CUT&RUN signal following nuclei purification.

We found that TIC-TaC coupled with CUT&Tag provides sufficient sensitivity and cell-type specificity to capture dynamic, activity-dependent chromatin remodeling within a genetically defined neuronal population in vivo. By profiling four histone PTMs in parallel in the same drug-induced seizure paradigm, we uncovered activity-dependent dynamics for both active and repressive marks. H3K27ac was the most tightly coupled to seizure-induced gene expression, showing concordant increases at upregulated genes and the strongest enrichment by GSEA, consistent with its established role as a marker of active promoters and enhancers.^31,66^ H3K27me3 showed a complementary but more limited relationship with transcription, decreasing at a subset of upregulated genes and increasing at non-differentially expressed which supports a model in which loss of Polycomb-mediated restrains contributes to activity dependent gene induction. Despite their association with active transcription, H3K9ac and H3K4me3 were largely uncoupled from seizure-induced transcriptional changes. This suggests that these marks may reflect a more permissive chromatin state rather than directly driving transcriptional output, or that their dynamics change on a different timescale than captured here.

We demonstrated that neuronal activity remodels the histone variant landscape in excitatory neurons following seizure. We observed a global loss of H2BE, consistent with prior work demonstrating that H2BE is inversely correlated with neuronal activity.^28^ Because H2BE promotes chromatin accessibility, depletion of H2BE may act to reduce transcription of synaptic genes and reduce neuronal excitability following seizure. In support of this model, a large subset of downregulated genes related to ion channels, receptors and synaptic organization/adhesion lost H2BE upon seizure induced neuronal activity. H2A.Z was modestly increased following seizure induced neuronal activity suggesting a distinct response to other forms of stimuli such as fear conditioning.^65,67^ Differences in stimulus paradigm (seizure vs. fear conditioning), brain region (cortex vs. hippocampus), or cell type profiled (excitatory neurons vs. bulk tissue) may account for this divergence. However, we did recapitulate the reported H2A.Z decrease at *Egr1* and gain at *Ppp3ca* suggesting that locus specific activity induced dynamics are consistent.^65^

TIC-TaC represents an efficient method that enables cell-type specific epigenomic profiling of specific cell types in complex tissues. In this study, we specifically profiled excitatory neurons by combining TIC-TaC mice with Nex-Cre mice, which demonstrated the ability to perform both CUT&RUN and CUT&Tag profiling of multiple marks. Further, by using drug-induced seizures we found that we can profile dynamic changes in marks and histone variants, allowing us to probe how the epigenome changes as a function of neuronal activity. In total, TIC-TaC represents an adaptable method that will facilitate epigenomic profiling of a diverse set of cell and tissue types, helping to reveal novel biology.

## Limitations

In this study, we focused on the use of TIC-TaC to profile histone PTMs and histone variants in excitatory forebrain neurons. The TIC-TaC method requires a Cre-dependent mouse line to conduct cell-type specific profiling *in vivo*. If no cell-type specific Cre exists for a cell line of interest, this approach will not be appropriate. Further, while it is likely that this approach could also be used in combination with viral injections of Cre-expressing vectors to avoid additional transgenic lines, we have not tested this. We also recognize that the time of the protocol is significant. In our hands, it takes approximately 4 hours to complete all steps up to binding of primary antibodies for standard CUT&RUN/Tag workflows. Despite this time investment, TIC-TaC enables preparation of multiple samples in parallel which may ultimately be more efficient than other approaches such as cell sorting which must be performed sequentially. Further, we have thus far only tested a single Cre line but hope that this method will be broadly applicable beyond the neuronal system we used here. By creating a Cre-dependent system, TIC-TaC can be applied to numerous cell types in the brain or beyond, provided there is a cell-type specific Cre. This will enable epigenetic profiling of a broad range of cell types in many different tissues.

## Lead contact

Erica Korb

## Materials availability

Mice are available upon request.

## Data and code availability

This paper analyzes the following publicly available datasets: CUT&RUN from 2-month-old mouse prefrontal cortex (GSE222072),^40^ CUT&RUN from FACS-isolated excitatory neuronal nuclei (GSE218312),^41^ chromatin immunoprecipitation sequencing (ChIP-seq) from FACS-isolated neuronal nuclei (GSE246772), and mouse cortical single nucleus Drop-seq from PTZ or home cage mice (GSE280192).^45^

## Declaration of generative AI and AI-assisted technologies in the writing process

When preparing the manuscript, the authors used Claude to improve clarity and grammar after an initial draft was written. The authors reviewed and edited content after using this tool.

## Acknowledgements

We thank Dr. Chris Lengner and the Penn Vet Core for generation of the transgenic mice. This work was supported by the National Institutes of Health grant 1DP2MH129985 (EK), National Institutes of Health grant R01NS134755 (EK), National Institutes of Health grant R35 GM142505 (GMB), Autism Spectrum Program of Excellence at the University of Pennsylvania, Eagles Autism Foundation, the Brody Foundation Fellowship (BL), National Institutes of Health grant 1T32GM156697 (KS). We would also like to thank members of the Korb and Burslem labs for valuable discussions.

## Author contributions

Conceptualization: BL; methodology: BL, KS; investigation: BL, KS, AS; analysis: BL, KS; Resources: CC, JB, GMB; sequencing data analysis and visualization: KS, supervision: EK, GMB; writing: BL, KS, EK; funding acquisition: EK, GMB.

## Methods

### Cloning

#### Cloning of Sun1-TEV-Constructs

TEV sites were introduced into the pCDNA.3_Sun1-GFP vector (gift by the Jeremy Nathans lab at Johns Hopkins) using standard cloning practices. In short, primers designed to incorporate a TEV site (See Primers table) were used to amplify the vector backbone using Q5 DNA polymerase (NEB, cat no. M0491L) following manufacturer instructions. Following a DpnI digest Amplicons were directly transformed into NEBalpha competent cells and sequences were confirmed through plasmidsaurus long read nanopore sequencing.

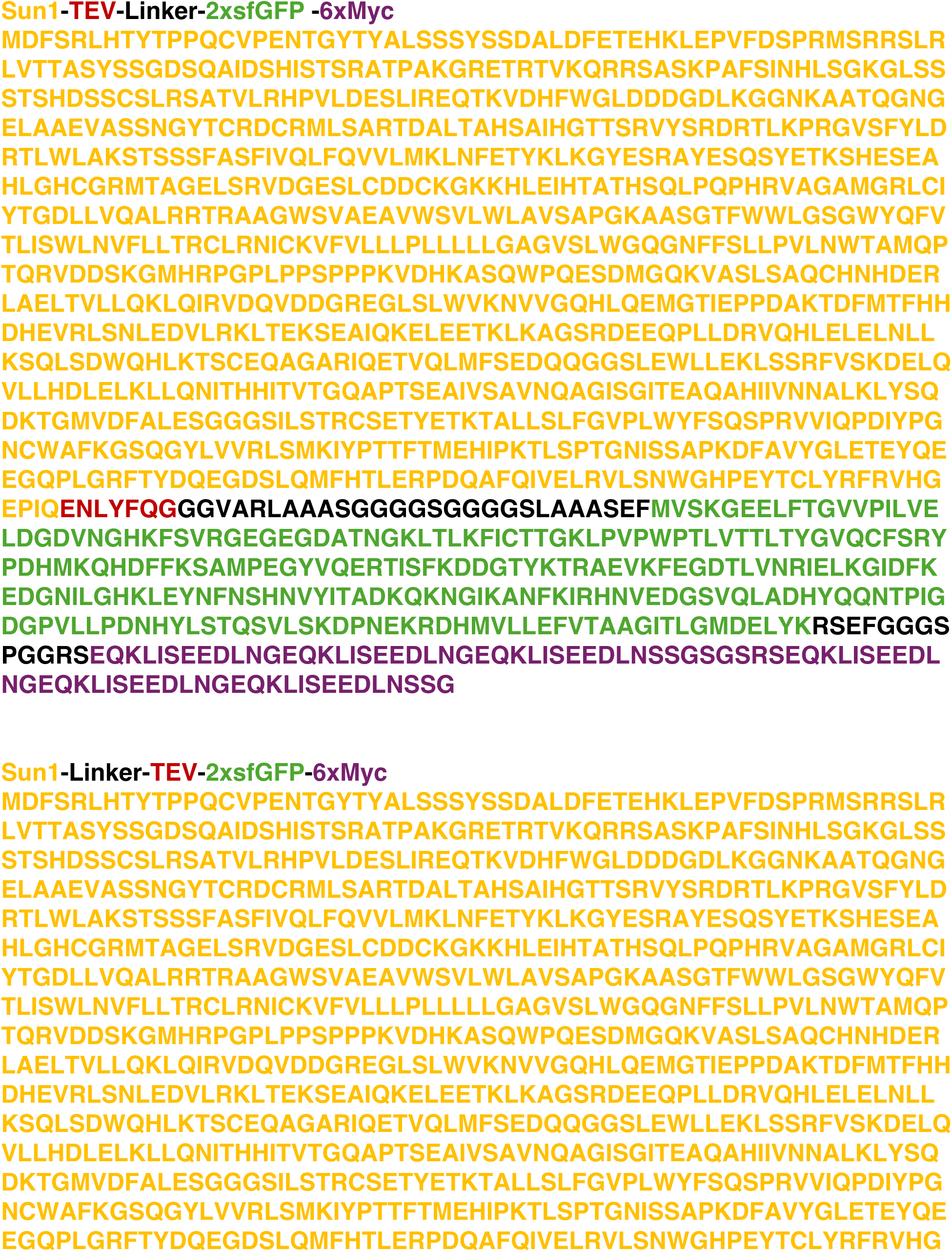
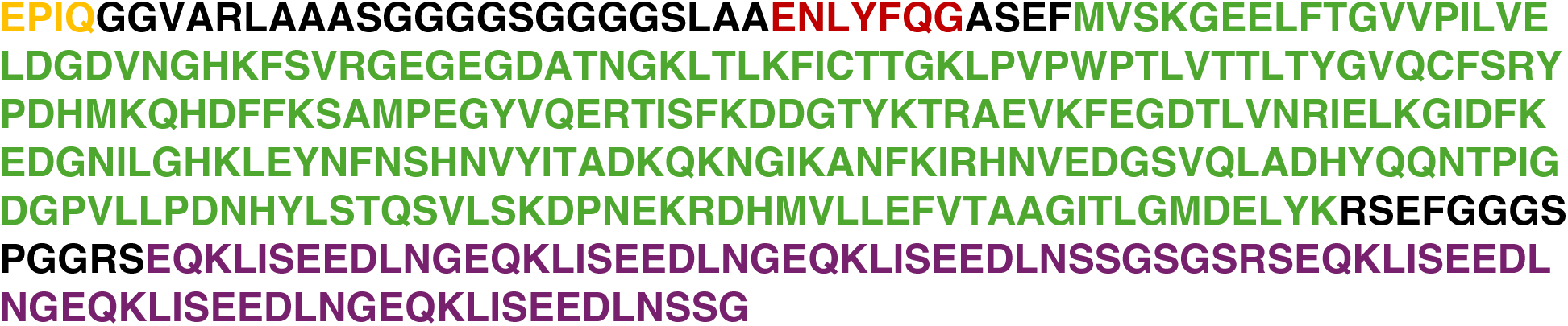

#### Cloning Rosa26 Targeting Construct

Sun1-TEV-GFP was subcloned into an in house Rosa26 targeting vector using a Gibson Assembly approach. In short, the targeting vector was digested with a combination of BstbI (NEB) and MluI-HF (NEB) in rCutsmart buffer for 1 hour at 37C. Sun1-TEV-GFP was amplified using primers with designed overhangs with the digested vector (Primers table) using Q5 DNA polymerase as described above. The fragments were then assembled using NEB HiFi Assembly Mastermix (NEB) following manufacturer instructions. Following assembly, the product was directly transformed into NEBalpha competent cells and sequences were confirmed through plasmidsaurus long read nanopore sequencing.

### Recombinant Protein Expression and Purification

#### TEV

pET-24d(+) containing His6-TEV protease (gifted by the Richard Baxter lab at Yale) was transformed into BL21(DE3) cells. Cells were grown in ZY-5052 auto-inducing media containing 50 μg/mL kanamycin at 37 °C while shaking at 200 RPM. At an OD600 of ∼1, the temperature of the incubator was changed to 20 °C and cells were left to shake overnight. Cell pellets were harvested and resuspended in lysis buffer (50 mM Tris-HCl pH 8.0, 300 mM NaCl, 20 mM imidazole, 20% glycerol v/v, 0.2% NP-40 v/v, 10 mM MgSO4) supplemented with EDTA-free protease inhibitor tablet (Thermo Scientific, cat. #A32955). Resuspended cells were sonicated for 15 min (2 sec on, 2 sec off, 40% amplitude) and spun down. Supernatant was incubated with Ni-NTA resin for 1 hr, shaking at 4 °C. Resin was washed with TEV binding buffer (50 mM Tris-HCl pH 8.0, 300 mM NaCl, 20 mM imidazole, 20% glycerol v/v) and elution buffer (50 mM Tris-HCl, pH 8.0, 300 mM NaCl, 300 mM imidazole, 20% glycerol v/v). Eluted protein was then dialyzed overnight into desalting buffer (50 mM Tris-HCl pH 8.0, 150 mM NaCl, 20% glycerol v/v). Dialyzed protein was quantified and stored in desalting buffer with 50% glycerol (v/v) at a 2 mg/mL concentration.

#### Open-Tn5

pG-OpenTn5 containing a His10 tag and mutations E54K, L372P was produced as described previously.^68^ The plasmid for pG-Tn5 was first transformed into BL21(DE3) cells. Cells were grown in TB supplemented with 50 μg/mL kanamycin and 0.4% glycerol while shaking at 37 °C. At an OD600 of ∼1.2, cells were induced with 0.6 mM IPTG and left shaking overnight at 18 °C. Cell pellets were harvested and resuspended in LysEQ buffer (20 mM HEPES pH 7.8, 800 mM NaCl, 20 mM imidazole, 10% glycerol, 1 mM EDTA, 2 mM TCEP-HCl) supplemented with EDTA-free cOmplete protease inhibitor (Roche, cat #11873580001). Resuspended cells were sonicated for 10 min (2 sec on, 2 sec off, 40% amplitude) and spun down. Lysates were run twice through Ni-NTA resin and washed with WashB1 buffer (20 mM HEPES pH 7.8, 800 mM NaCl, 30 mM imidazole, 10% glycerol, 1 mM EDTA, 2 mM TCEP-HCl), WashB2 buffer (20 mM HEPES pH 7.8, 800 mM NaCl, 45 mM imidazole, 10% glycerol, 1 mM EDTA, 2 mM TCEP-HCl), and EluB buffer (20 mM HEPES pH 7.8, 800 mM NaCl, 400 mM imidazole, 10% glycerol, 1 mM EDTA, 2 mM TCEP-HCl). Fractions containing protein were dialyzed overnight into SECB buffer (50 mM Tris-HCl pH 7.5, 800 mM NaCl, 10% glycerol, 0.2 mM EDTA, 2 mM DTT). Dialyzed protein was run on a HiLoad 16/600 Superdex 200 column (Cytiva, product #28989335) in a 1.5 column volume isocratic elution at a flow rate of 1 mL/min. Fractions containing protein were concentrated, quantified and stored in SECB buffer with 55% glycerol (v/v) at a 1 mg/mL concentration.

### Cell Culture

#### Cell Lines

HEK293 cells cell lines were obtained from ATCC and routinely tested for mycoplasma. Cells are maintained at a low (<20) passage number.

#### Transfections

HEK293 cells in a 12-well format were transfected using Lipofectamine 2000 following manufacturer instructions. In short, when cells reached ∼ 60% confluency a total of 2.1 µL of Lipofectamine 200 was diluted in 52.5 µL of Optimem in a 1.5 mL tube. In another tube, 2.4 µg of DNA (TIC-TaC constructs TEVL and LTEV) were diluted in 52.5 µL of Optimem. Each of these solutions were incubated for 5 minutes at room temperature before mixing and incubating for a minimum of 15 minutes at room temperature. After incubating, 100 µL of the lipofectamine and DNA mixture was added dropwise to the cells. The following day, media was removed and replaced with fresh. Cells were harvested 2 to 3 days after transfection, or once GFP expression was clearly observed in at least half the cells.

### Mice

#### Breeding and Housing

TIC-TaC mice were generated in this study and were crossed to Nex-Cre mice to obtain double heterozygous and double homozygous (TIC-TaC;Nex-Cre) mice used in this study. Genotyping was performed for both as described below. Mice were housed in a 12-hour light-dark cycle and fed a standard diet. Experiments were performed in accordance with and approval of IACUC.

#### Generation of TIC-TaC Mouse

To generate the TIC-TaC mouse, the Rosa26 targeting vector was first linearized via restriction digest with AclI (NEB) for 1 hour at 37C. The Transgenic Mouse Core (Penn Vet, University of Pennsylvania) performed pronuclear injection to generate generate TIC-TaC +/-embryos. In brief, the linearized targeting vector was introduced by nucleofection (Amaxa Neucleofactor System, Lonza) into V6.5 embryonic stem cells, after which the cultures were subjected to G418 selection for 7 days and drug-resistant colonies were subcloned and screened for proper insertion. Several correctly targeted clones were identified, and one was injected into C57Bl/6 blastocysts, resulting in the generation of several high-contribution male chimeras. Chimeric founder mice were crossed to C57Bl/6J female mice and progeny were screening for successful transgene transmission using a primer set with a common forward primer that binds in the left Rosa26 homology arm, and WT and Mutant reverse primers that bind within the Rosa26 locus and CMV promoter, respectively.

#### Genotyping

Ear punches collected from P14 pups were boiled in 50 μL of denaturing solution (25 mM NaOH, 0.2 mM EDTA) for 1 hour. Solutions were then neutralized with 40 mM Tris.HCl pH 4.0. Genomic DNA was purified using a Zymo DNA Clean and Concentrate kit using manufacturer instructions. Genotyping PCR was performed using KAPA2G () using manufacturer instructions (see primer tables).

### Seizure Induction

Pentylenetetrazol (PTZ; Sigma P6500) was injected intraperitoneally at 50mg/kg (in PBS). Control mice received PBS injections at equivalent volume. Mice were observed for 15 minutes after injection to score seizures, then sacrificed. The modified Racine scale was used to measure seizure induction as follows.

- Stage 1: Hypoactivity culminating in behavioral arrest with contact between abdomen and the cage.
- Stage 2: Partial clonus (PC) involving the face head or forelimbs.
- Stage 3: Generalized clonus (GC) including all four limbs and tail, rearing or falling.
- Stage 4: Generalized Tonic-Clonic seizure (GTC)

Seizure susceptibility score was calculated as: (0.2)(1/PC latency) + (0.3)(1/GC latency) + (0.5)(1/GTC latency).

### Antibodies

Rabbit anti-GFP: ThermoFisher G10362 (RRID:AB_2536526)

GFP-Trap Magnetic Particles: Proteintech gtdk-20 (RRID:AB_2827592)

Rabbit anti-H3K27ac: Active Motif, 39133 (RRID:AB_2561016)

Rabbit anti-H3K9ac: Active Motif, 39917 (RRID:AB_2616593)

Rabbit anti-H3K4me3: Active Motif, 39159 (RRID: AB_2615077)

Rabbit anti-H3K27me3: Active Motif, 39156 (RRID:AB_2636821)

Rabbit anti-H2BE: Millipore ABE1384 (RRID:AB_3106370)

Rabbit anti-H2A.Z: Abcam ab4174 (RRID:AB_304345)

Normal Rabbit IgG: Sino Biological CR1 (RRID:AB_3073921)

Guinea Pig anti-Rabbit IgG: Antibodies-online ABIN101961 (RRID: AB_10775589)

### TEV Induced Cleavage of Nuclei Tagged in Specific Cell Types (TIC-TaC)

#### Gradient Isolation of Nuclei

Snap frozen brain tissue from TIC-TaC^+/-^; Nex-Cre^+/-^mice or frozen cell pellets from transfected HEK293s were homogenized in 1.2 mL of cold Buffer HB (20 mM Tricine-NaOH pH 7.8, 25 mM KCl, 1 mM dithiothreitol (DTT), 0.15 mM spermine, 0.5 mM spermidine, supplemented with protease inhibitor (Roche04693124001)) in a Dounce homogenizer. Tissue was broken up with 20 pulses of the loose pestle followed by 10 pulses of the tight pestle. To this, 107 µL of 5% Wash Buffer (HB supplemented with 5% IGEPAL-CA630) was added and samples were pulsed 7 more times with the tight pestle. Crude lysates were diluted two-fold with working solution (50% iodixanol, 20 mM Tricine-NaOH pH 7.8, 25 mM KCl, 0.17 mM DTT, 25 uM spermine, 83 uM spermidine) to yield a final concentration of 25% iodixanol. A gradient was prepared in a 15 mL falcon tube by placing 1.33 mL of 40% iodixanol and overlaying it with 2.5 mL of 30% iodixanol. Iodixanol concentrations are obtained by diluting working solution with HB. The crude lysate was then carefully placed on top of the 30% gradient and tubes were sealed. Nuclei were isolated by centrifugation (3200 RCF, 20 minutes, max acceleration, deceleration of 1, 4C). Following centrifugation, 1.2 mL was collected from the nuclei containing interface of the 30% and 40% layers. Nuclei were quantified using a hemacytometer. For TRAP samples, GFP positive nuclei were purified via immunoprecipitation as described below.

#### Isolation of Nuclei Tagged in Specific Cell Types (INTACT)

GFP positive nuclei were purified using INTACT as previously described with minor modifications. In short, 80 µL of ProteinA Dynabeads (ThermoFisher 10002D) was equilibrated with Wash Buffer (20 mM Tricine-NaOH pH 7.8, 25 mM KCl, 1 mM dithiothreitol (DTT), 0.15 mM spermine, 0.5 mM spermidine, 0.4% IGEPAL-CA630, supplemented with protease inhibitor) by washing twice with 400 µL. Samples were cleared of non-specific interactors by adding 20 µL of washed beads to isolated nuclei. Samples were then rotated end-to-end for 10 min at 4C. Following the pre-clear step, samples were placed on a magnetic rack for 3 minutes, and the supernatant was transferred to a clean tube containing 160 µL of wash buffer and 10 µL of rabbit a-GFP. Samples were then incubated at 4C with end-to-end rotation for 30 minutes. After antibody binding, the remaining 60 µL of equilibrated ProteinA Dynabeads were added and incubated for 30 minutes at 4C with end-to-end rotation. Samples were placed on a magnetic rack for 1 minute and then were resuspended by inverting. This step was repeated 8 times. On the last repeat, samples were placed on the rack for 3 min, and the supernatant was removed. Beads were washed 4 times with 400 µL of Wash Buffer. During each wash, samples were gently resuspended by inversion before placing back onto the rack. Following washing, nuclei were analyzed for purity using a hemocytometer before proceeding to TEV cleavage.

#### TEV Induced Cleavage

Following counting, bead bound nuclei were diluted or resuspended to a concentration of 1.25 million nuclei per mL. For cleavage of 200,000 nuclei, 160 µL of bead bound nuclei were collected and placed into a new tube. To each sample, 10 µL of 20x TEV buffer (1M Tris-HCl pH 8.0, 10 mM EDTA), 4 µL of 100 mM DTT, 13.4 µL of TEV (1:15 dilution), and 12.6 µL of wash buffer was added. Samples were mixed by pipetting and incubated for 2 hours at room temperature with rotation (3 hours at 4C for cultured cells transfected with Sun1-TEV-GFP). After incubation, samples were placed on a magnetic rack until clear and the beads were then resuspended via pipetting after clearing. Following resuspension, samples were returned to the magnetic rack, and once clear the supernatant containing nuclei was collected. For experiments comparing nuclei with or without TEV cleavage, TEV protease was substituted with wash buffer for -TEV samples. CUT&Tag and CUT&RUN workflows using approximately 100,000 nuclei were immediately started following TEV cleavage.

#### Blocking of pA-Dynabeads with IgG

Following nuclei purification, pA-Dynabeads were incubated with a saturating amount of IgG (determined using 30 µg binding capacity per 100 µL of pA-Dynabeads) in wash buffer. Bead-bound nuclei were incubated with IgG for 1 hour at 4C. IgG’s were selected based on the species that primary antibodies for targets of interest were raised in to avoid non-specific binding of subsequent secondary antibodies. For antibodies raised in Rabbit, a Ms-IgG was used to block, and for antibodies raised in Mouse, a Rb-IgG was used. Following incubation, samples were placed on a magnetic rack beads were washed 3 times with 400 uL of wash buffer. Samples were then used CUT&Tag or for direct staining of isolated nuclei.

### GFP nanobody nuclei immunoprecipitation

GFP-nanobody immunoprecipitation was performed as previously published.^29^ Snap frozen brain tissue from TIC-TaC^+/-^; Nex-Cre^+/-^mice was homogenized with a Dounce homogenizer in 1.5 mL of low sucrose buffer (15 mM Tris-HCl pH 7.4, 25 mM KCl, 5 mM MgCl2, 1 mM DTT, EDTA-free Protease Inhibitor, 0.5 mM Spermidine, 0.15 mM Spermine, 250 mM Sucrose). After homogenization, 1.5 µL of NP-40 was added to achieve a concentration of 0.1%. Homogenates were passed through a 70 µm filter, followed by a 40 µm filter, then centrifuged at 600 RCF at 4°C for 10 minutes. 900 µL of sucrose cushion buffer (15 mM Tris-HCl pH 7.4, 25 mM KCl, 5 mM MgCl2, 1 mM DTT, EDTA-free protease inhibitor, 0.5 mM Spermidine, 0.15 mM Spermine, 1.2 M Sucrose) was added to the suspension and gently mixed, then overlaid on top of 500 µL of sucrose cushion buffer. The mixture was centrifuged at 3,200 RCF at 4°C for 45 minutes. Following centrifugation, the supernatant was discarded and the pellet was resuspended in 1 mL of wash buffer (15 mM Tris-HCl pH 7.4, 25 mM KCl, 5 mM MgCl2, 1 mM DTT, EDTA-free Protease Inhibitor, 0.5 mM Spermidine, 0.15 mM Spermine, 250 mM Sucrose, 0.1% NP-40). Nuclei were counted and split into aliquots of 500,000. GFP-Trap Magnetic Particles (Proteintech, gtdk-20) were equilibrated and washed in wash buffer, then 25 µL was added to each 500,000 nuclei aliquot. The mixture was incubated at 4°C for 1 hour with rotation. After incubation, the magnetic particles were washed 3 times in 500 µL of wash buffer and nuclei were counted. INTACT nuclear isolation was performed in parallel to the sucrose cushion/GFP-trap method using the same brain tissue. The sucrose cushion/GFP-trap workflow was additionally performed using dilutions of 500,000, 200,000, 100,000, and 50,000 nuclei to test if nuclei number influenced clumping.

### Staining and Imaging

#### Perfusions

Animals were anesthetized in a chamber with a constant flow of Oxygen and Isoflurane. Following confirmation of anesthetization via a toe pinch, animals were fixed with a nose cone for transcardial perfusions. In short, after cutting the right atrium, animals were injected with 10-15mL of PBS to wash out blood, followed by administration of 15-20mL of formalin to fix all tissues. Following fixation, the whole brain was carefully extracted and placed in formalin to incubate overnight at 4°C. Following overnight fixation, brains were transferred to 30% sucrose for 48 hours at 4°C. Each brain was then embedded in OCT and was stored at -80°C until sectioning.

#### Immunohistochemical Staining

OCT embedded brains were sectioned using a cryostat to collect 16µm slices onto SuperFrost Plus slides (Fisher #22-037-246). Immediately following sectioning, each slide was dried completely at room temperature before washing in TBS to remove excess OCT. Sections were blocked for 1 hour at room temperature in blocking solution (3% BSA, 0.1% Triton in TBS). GFP and NeuN Staining were performed by preparing 1:500 dilutions of rabbit anti-GFP-488 (Thermo #A-21311) and mouse anti-NeuN-555 (Sigma #MAB377A5) in blocking solution and incubating for 1 hour at room temperature. Following incubation, sections were washed once with TBS by incubating for 5 minutes. Following first wash each section was incubated with 1:5000 DAPI in TBS for 5 minutes. Following DAPI incubation, each section was wash twice by incubating with TBS for 5 minutes. After the final wash, sections were allowed to dry briefly before adding 10µL of Prolong Gold solution over each tissue section. A coverslip was gently placed over top of the slide and was then dried overnight in the dark.

#### Staining of Isolated Nuclei

For indicated samples, pA-Dynabeads were blocked with excess Rb-IgG as described above. Nuclei were incubated overnight at 4C with a Ms-H3K27ac antibody (1:100, in 100 µL of wash buffer). Following overnight incubation, beads were washed twice with 400 µL of wash buffer. Following washes, nuclei were incubated with Ms-AF555 (1:100 in 100 µL of wash buffer) for 1 hour at 4C. After incubation, beads were washed twice with 400 µL of wash buffer. After the final wash, nuclei were resuspended in 400 uL of wash buffer. with 1:5000 diluted DAPI.

#### Imaging of Isolated and Purified Nuclei

All quantification and imaging of isolated or purified nuclei was performed on disposable hemocytometers. Prior to loading, nuclei were diluted ten-fold (crude isolated nuclei) or two-fold (purified nuclei) in wash buffer containing 1:5000 DAPI. A disposable hemocytometer was loaded with 10 uL of sample and images were taken on a (insert scope) fitted with a (laser and filter apparatus).

#### Imaging of TIC-TaC Tissue

Stained TIC-TaC slices were imaged using a Leica DM 6000, TCS SP8 microscope and a ×20 HC PL APO CS2 oil objective with an NA of 1.40. Images were taken of the hippocampus and cortex. Processing was performed using ImageJ for qualitative comparisons of TIC-TaC^+/-^; Nex-Cre^+/-^ and TIC-TaC^+/-^; Nex-Cre^-/-^tissue.

### CUT&Tag-sequencing

CUT&Tag was performed as previously published.^20^ Concanavalin A-coated beads (EpiCypher, 21-1411) were washed two times with 1 mL of filter-sterilized binding buffer (20 mM HEPES pH 7.9, 10 mM KCl, 1 mM CaCl2, 1 mM MnCl2) and resuspended in 11 µL per reaction of binding buffer. 11 µL of activated conA beads were added to 100,000 nuclei isolated by TIC-TAC, then washed once with 100 µL of antibody buffer (20 mM HEPES pH 7.5, 150 mM NaCl, 0.5 mM Spermidine, EDTA-free Protease Inhibitor, 0.05% Digitonin, 2 mM EDTA, 1% BSA). For histone modification seizure experiments, 1% of the input nuclei number of HEK293 cells were added to TIC-TAC isolated nuclei as a spike-in control. Nuclei bound conA beads were resuspended in 50 µL of antibody buffer and 1 µg of primary antibody was added to each reaction: Rabbit anti-H3K27ac (Active Motif, 39133), Rabbit anti-H3K9ac (Active Motif, 39917), Rabbit anti-H3K4me3 (Active Motif, 39159), Rabbit anti-H3K27me3 (Active Motif, 39156), Rabbit anti-H2BE (Millipore ABE1384), Rabbit anti-H2A.Z (Abcam ab4174), Normal Rabbit IgG (Sino Biological, CR1). Samples were vortexed, then incubated overnight at 4°C on a rocking shaker. Beads were collected on a magnetic rack, the supernatant was discarded, and 50 µL of secondary antibody (Guinea Pig anti-Rabbit IgG: Antibodies-online ABIN101961) diluted 1:100 in Digi-Wash buffer (20 mM HEPES pH 7.5, 150 mM NaCl, 0.5% Spermidine, EDTA-free protease inhibitor, 0.05% digitonin) was added. Samples were subsequently incubated at room temperature for 1 hour on a nutator, then washed twice with Digi-Wash buffer. Beads were collected on a magnet, supernatant removed, and beads were resuspended in 50 µL of loaded pG-Tn5 adapter complex (diluted 1:100 in Digi-300 Buffer; 20 mM HEPES pH 7.5, 300 mM NaCl, 0.5 mM spermidine, 0.01% digitonin).^68^ Samples were incubated at room temperature for 1 hour on a nutator, then washed twice with Digi-300 buffer. Beads were then collected on a magnet, supernatant removed and resuspended in 300 µL of Tagmentation Buffer (20mM HEPES pH 7.5, 300 mM NaCl, 0.5 mM spermidine, 0.01% digitonin, 10 mM MgCl2). Samples were incubated for 1 hour at 37°C in a heat block. Tagmentation was stopped by adding 10 µL 500 mM EDTA, 3 µL 10% SDS, and 2.5 µL of Proteinase K (20 mg/mL). Protein was digested by incubating samples for 1 hour at 55°C in a heat block, then DNA was isolated using the DNA Clean and Concentrator-5 Kit (Zymo Research D4014). A universal i5 primer and a uniquely barcoded i7 primer were ligated, and libraries were amplified by PCR using NEBNext High-Fidelity 2X PCR Master Mix (NEB M0541). The following cycling conditions were used: 72°C for 5 minutes, 98°C for 30 seconds, 98°C for 10 seconds & 63°C for 10 seconds (14 cycles), 72C for 1 minute, 8°C hold. Library clean-up was performed using a 1.3X ratio of AMPure XP beads (Beckman Coulter, A63881) and eluted off beads in 25 µL of Tris-HCl, pH 8. Prior to sequencing, library size distribution and quality was assessed by capillary electrophoresis using an Agilent 4200 TapeStation with high sensitivity D1000 reagents (Agilent, 5067-5585, 5067-5584). Libraries were quantified by qPCR using a KAPA Library Quantification Kit (Roche, 07960140001). Libraries were sequenced on an Illumina NextSeq 1000 instrument (61-bp read length, paired-end, with dual 8-bp indices).

### CUT&RUN-sequencing

CUT&RUN was performed as previously published with modifications.^19^ Concanavalin A-coated beads (EpiCypher, 21-1411) were washed two times with 1 mL of Binding Buffer and resuspended in 11 µL per reaction of Binding Buffer. 11 µL of activated conA beads were added to 100,000 nuclei isolated by TIC-TAC, then washed once with 100 µL of antibody buffer. Nuclei bound conA beads were resuspended in 50 µL of Antibody Buffer and 1 µg of primary antibody was added to each reaction: Rabbit anti-H3K27ac (Active Motif, 39133), Rabbit anti-H3K9ac (Active Motif, 39917), Rabbit anti-H3K4me3 (Active Motif, 39159), Rabbit anti-H3K27me3 (Active Motif, 39156), Normal Rabbit IgG (Sino Biological, CR1). Samples were vortexed, then incubated overnight at 4°C on a rocking shaker. Beads were collected on a magnetic rack, supernatant was discarded, and beads were washed twice with Digitonin Buffer (20 mM HEPES pH 7.5, 150 mM NaCl, 0.5 mM Spermidine, 0.1% Digitonin, EDTA-free protease inhibitor). Beads were resuspended in 50 µL of Digitonin Buffer, and 2.5 µL of pAG-MNase (EpiCypher, 15-1116) was added. Samples were gently vortexed to mix and incubated for 10 minutes at room temperature on a nutator. Beads were collected on a magnetic rack, supernatant was discarded, and beads were washed twice with Digitonin Buffer. Beads were resuspended in 50 µL of Digitonin Buffer, 1 µL of 100 mM CaCl2 was added, and samples were incubated for 30 minutes on wet ice. To stop MNase cleavage, 33 µL of Stop Buffer was added (340 mM NaCl, 20 mM EDTA, 4 mM EGTA, 50 µg/mL RNAse A, 50 µg/mL Glycogen). Samples were incubated for 10 minutes at 37°C in a thermocycler. Following incubation, samples were placed on a magnet and supernatant containing CUT&RUN enriched DNA was collected. DNA was purified from the supernatant using a 1.4X ratio of CUTANA DNA purification beads (EpiCypher, 21-1407-50ML). Library preparation was performed using the CUTANA CUT&RUN Library Prep Kit (EpiCypher, 14-1001 & 14-1002). End repair was performed by adding 5 µL of End Repair Master Mix (End Prep Buffer & End Prep Enzyme) to each sample and run in a thermocycler with a heated lid (75C) with the following program: 20°C for 20 minutes, 65°C to 30 minutes, hold at 4-12°C. Adapter ligation and U-excision was carried out by adding 1.25 µL of 1.5 µM Adapter for Illumina and 15.5 µL of Ligation Master Mix (Ligation Mix & Ligation Enhancer) to end repaired DNA and incubating for 15 minutes at 20°C. Next, 1 µL of U-Excision Enzyme was added to each reaction and incubated at 37°C in a thermocycler (lid temperature 47°C) for 15 minutes. DNA cleanup was performed using 1X ratio of CUTANA DNA purification beads and DNA was eluted in 12 µL of 0.1X TE Buffer. 1 µL of uniquely barcoded i7 primer, 1 µL of uniquely barcoded i5 primer, and 12.5 µL of Hot Start 2X PCR Master Mix were added to each reaction and vortexed to mix. Reactions were placed in a thermocycler with the lid set to 105°C and the following program was run: 98°C for 45 seconds, 98°C for 15 seconds & 60°C for 10 seconds (14 cycles), then 72°C for 60 seconds, 4-12°C hold. Libraries were cleaned up using a 1X ratio of CUTANA DNA purification beads and eluted in 12 µL of 0.1X TE Buffer. Prior to sequencing, library size distribution and quality was assessed by capillary electrophoresis using an Agilent 4200 TapeStation with high sensitivity D1000 reagents (Agilent, 5067-5585, 5067-5584). Libraries were quantified by qPCR using a KAPA Library Quantification Kit (Roche, 07960140001). Libraries were sequenced on an Illumina NextSeq 1000 instrument (61-bp read length, paired-end, with dual 8-bp indices).

### CUT&Tag/RUN data processing and analysis

#### Pre-processing and normalization

Raw BCL data were converted to FASTQ format and demultiplexed using BCL Convert (v4.3.13). Reads were aligned to the Mus musculus genome build mm10 with Bowtie2 (v2.4.5). Biological replicates (n=3) were subsampled to equivalent read depth per target using samtools (v1.22.1). TIC-TAC CUT&Tag samples without TEV (Figure 2) had significantly fewer reads compared to samples treated with TEV, thus we did not subsample each sample to the same number of reads. TIC-TaC CUT&RUN replicates were subsampled to the same number of reads per target. For benchmarking experiments in Figure 4, in the case that 2 biological replicates were used in publicly available datasets, we subsampled reads such that the bigWigs merged by condition had equivalent read depth. For seizure experiments in Figure 5, n=2 (home cage) and n=3 (seizure) replicates were used; we subsampled reads such that bigWigs merged by condition had equivalent read depth. Reads across replicates were merged using samtools and bamCoverage from deepTools (v3.5.6) was used to create bigWigs from merged BAM files.

#### Spike-in normalization

For histone modification seizure experiments, reads were aligned separately to the experimental genome (Mus musculus build mm10) and the spike-in genome (Homo sapiens build hg38) with Bowtie2. The fraction of spike-in reads relative to experimental reads was calculated, and the sample with the smallest spike-in fraction was used as the reference. Normalization factors for all other samples were calculated to the reference such that all scaling factors were less than 1. Normalized bigWigs were generated using bamCoverage (scaleFactor). Raw read counts for specific genomic regions were scaled using normalization factors in RStudio.

#### Visualization

Merged bigWigs were plotted as heatmaps with metaplots displaying average signal profiles using BED files containing gene body coordinates for either neuronally expressed genes or lowly/non-expressed neuronal genes. Gene expression level in WT primary neurons was previously published.^45^ Genes with base mean less than 3 were defined as non-expressed and the remaining genes were divided into 3 terciles and categorized as low-mid- and high expressed. Neuronally expressed genes included low, mid, and high expression categories. An additional 2 or 4 kb were plotted on either end of the TSS and TES. CUT&Tag normalized reads were plotted based on expression bins and the Wilcoxon rank-sum test was used to compare between seizure and home cage conditions. Genome browser tracks from merged bigWigs were created in IGV (v2.19.17).

#### Correlation analysis

Merged BAM files were used to perform quantitative correlation analysis. For H3K27ac, H3K9ac, and H3K4me3 BED files with TSS coordinates plus and minus 500bp were used. For H3K27me3 genic correlations, a BED file with gene body coordinates of lowly or non-expressed neuronal genes (minus 500bp at the TSS and TES) was used. For H3K37me3 TSS correlations, a BED file with coordinates of TSSs at low-non expressed neuronal genes (plus and minus 500bp) was used. Spearman rank correlations were performed in RStudio (v2026.04.0+526).

#### Peak calling

Peaks were called with SEACR (v1.3)^69^ using stringent mode for H3K27ac, H3K9ac, H3K4me3, and relaxed mode for H3K27me3, normalized to IgG and annotated with HOMER (v5.1). Relaxed mode was used to capture broader H3K27me3 domains. A peak cutoff score of 10 was applied. Unique genes with peaks overlapped and visualized using https://bioinformatics.psb.ugent.be/webtools/Venn/.

### Analysis of snRNA-Drop sequencing

Cortical single nucleus Drop-seq from WT PTZ treated, or home cage mice was analyzed as reported previously.^45^ We selected the excitatory neuron clusters from seizure and control groups to match TIC-TAC isolated excitatory nuclei. Differential expression analysis was performed using the MAST framework by the FindMarkers function in Seurat (v5.4.0).^70^ The excitatory neuron clusters contained 9,949 and 10,830 nuclei for home cage and seizure groups, respectively. Genes were tested if detected in at least 10% of nuclei in either comparison group. Genes were considered differentially expressed at Bonferroni-corrected P<0.00001 and an absolute fold change threshold of 1.5. Pseudobulk analysis was performed with the AggregateExpression function in Seurat to merge counts across nuclei by condition. Differential expression analysis was performed on the pseudobulk count matrix using DEseq2 (v1.46.0). Genes were considered differentially expressed at a Benjamini-Hochberg adjusted p<0.05 and an absolute fold-change threshold of 1.5. To integrate CUT&Tag with snRNA-seq, we examined TSSs of differentially expressed genes (170 down, 100 up). In cases when multiple start sites were detected for these DEGs, we selected the site with the highest H3K4me3 CUT&Tag signal in the home cage control condition. Of the DEGs, 158 downregulated, 81 upregulated and 1,016 non differentially expressed had detectable CUT&Tag signal. CUT&Tag log2 fold change (seizure/home cage) at TSSs was plotted using the R package ComplexHeatmap (v2.22.0, Gu et al., 2016, PMID: 27207943). Concordance was defined as downregulated genes exhibiting a 1.2-fold increase in H3K27me3, 1.2-fold increase in H2A.Z, and a 1.2-fold decrease in H3K27ac. For upregulated genes concordance was defined 1.2-fold decrease in H3K27me3, 1.2-fold decrease in H2A.Z or a 1.2-fold increase in H3K27ac. GSEA was performed in R using ClusterProfiler (v4.14.6).^71^ For each histone PTM, ranked lists were created ordered by log2 fold change (seizure/home cage). Upregulated and downregulated gene sets were tested for enrichment for each histone PTM ranked list, with significant enrichment at a threshold of padj <0.05.

---

**Supplementary Figure 1. TIC-TaC principle and validation. A)** Fluorescence imaging of pre-INTACT and post-INTACT purified nuclei from HEK293 cells transfected with pCDNA.3_Sun1-GFP, GFP (green) and DAPI (blue). **B)** Quantification of pre-INTACT and post-INTACT purified nuclei from HEK293 cells transfected with pCDNA.3_Sun1-GFP demonstrating enrichment for GPF positive nuclei. (*, p < 0.05). **C)** Schematic depicting challenges of combining INTACT with CUT&Tag. Nuclei bound pA-Dynabeads will result in antibodies binding to pA coating beads rather than intended epitopes. pA-Tn5 will then bind to antibodies on the Dynabeads instead of to any antibodies that reach the intended epitopes. **D)** Immunofluorescence of nuclei bound to pA-dynabeads that are not treated with additional antibodies, that are directly treated with an H3K27ac antibody, and that are blocked with excess Rb-IgG prior to the addition of an H3K27ac antibody. H3K27ac visualized by addition of a AlexaFluor-555 (red), with GFP (green), and DAPI (blue). **E)** Tapestation of CUT&Tag performed with an IgG blocking step in bead bound HEK293 GFP positive nuclei. As a positive control, isolated nuclei prior to INTACT were bound to ConA beads. Prior to CUT&Tag on INTACT purified nuclei, Dynabeads were blocked with either excess Rb-IgG or Ms-IgG. For CUT&Tag using Rb-IgG or Rb-H3K27ac, beads were blocked with excess Ms-IgG. For CUT&Tag using Ms-IgG or Ms-H3K27ac, beads were blocked with excess Rb-IgG. **F)** Fluorescence imaging of pre-INTACT and post-INTACT purified nuclei from TIC-TaC^+/-^; Nex-Cre^+/-^mice using GFP-nanobody beads demonstrating poor purity. **G,H)** Quantification of fluorescence imaging of pre-INTACT and post-INTACT purified nuclei from TIC-TaC^+/-^; Nex-Cre^+/-^mice using GFP-nanobody beads demonstrating poor purity compared to INTACT (G, ***, p<0.001) and that purity is not improved based on the number of GFP positive nuclei in solution prior to purification (H). **I)** Fluorescence imaging of pre-INTACT and post-INTACT purified nuclei from TIC-TaC^+/-^; Nex-Cre^+/-^mice using GFP-nanobody beads demonstrating clumping of nuclei. **J,K)** Schematic of two different designs for the TIC-TaC system. The TEV site was placed before the linker (J) or after the linker (K) that separates Sun1 from GFP. **L)** Fluorescence imaging of TEV cleavage time course comparing LTEV and TEVL constructs. Time course was conducted at 4C over the course of 3 hours. A minus TEV sample was used as the 0-hour time point. **M)** Quantification of the TEV cleavage time course comparing LTEV (dark red) and TEVL (light red). Near quantitative release was achieved within two hours and three hours respectively. **N)** Quantification of TEV cleavage time course directly comparing LTEV (dark red) and TEV (light red) at 1 hour and 2 hours which demonstrates that the LTEV configuration results in quicker elution from beads (*, p<0.05; ***, p<0.001).

**Supplementary Figure 2. TIC-TaC CUT&Tag/RUN cell-type-specific loci. A)** Representative CUT&Tag gene tracks for a neuronal marker gene (*Syt1*, left) and non-neuronal marker genes (astrocyte: *Aldh1l1*, middle; microglia: *Fcrls*, right). Tracks show n=3 merged biological replicates. **B)** Representative CUT&RUN gene tracks for a neuronal marker gene (*Syt1*, left) and non-neuronal marker genes (astrocyte: *Aldh1l1*, middle; microglia: *Fcrls*, right). Tracks show n=3 merged biological replicates.

**Supplementary Figure 3. TIC-TaC CUT&Tag/RUN extended benchmarking. A)** Correlation between H3K27ac TSS reads from TIC-TAC CUT&Tag versus TIC-TAC CUT&RUN (Spearman r=0.94, n=29,679), H3K27me3 gene body reads (Spearman r=0.88, n=6,650), H3K27me3 TSS reads (Spearman r=0.84, n=7,084), and H3K4me3 TSS reads (Spearman r=0.91, n=29,679). **B)** Correlation between H3K27ac TSS reads from TIC-TAC CUT&RUN versus bulk cortex CUT&RUN (Spearman r=0.89, n=29,679), versus FACS excitatory nuclei CUT&RUN (Spearman r=0.88, n=29,679), and versus FACS neuronal nuclei ChIP-seq (Spearman r=0.87, n=29,679). **C)** Correlation between H3K27me3 gene body reads from TIC-TAC CUT&RUN versus bulk cortex CUT&RUN (Spearman r=0.89, n=6,243), versus FACS excitatory nuclei CUT&RUN (Spearman r=0.86, n=6,243), and versus FACS neuronal nuclei ChIP-seq (Spearman r=0.78, n=6,243). **D)** Correlation between H3K27me3 TSS reads from TIC-TAC CUT&Tag versus bulk cortex CUT&RUN (Spearman r=0.86, n=7,084), versus FACS excitatory nuclei CUT&RUN (Spearman r=0.86, n=7,084), and versus FACS neuronal nuclei ChIP-seq (Spearman r=0.78, n=7,084). **E)** Correlation between H3K27me3 TSS reads from TIC-TAC CUT&RUN versus bulk cortex CUT&RUN (Spearman r=0.85, n=7,084), versus FACS excitatory nuclei CUT&RUN (Spearman r=0.83, n=7,084), and versus FACS neuronal nuclei ChIP-seq (Spearman r=0.77, n=7,084). **F)** Correlation between H3K4me3 TSS reads from TIC-TAC CUT&RUN versus bulk cortex CUT&RUN (Spearman r=0.92, n=29,679) and versus FACS excitatory nuclei CUT&RUN (Spearman r=0.90, n=29,679).

**Supplementary Figure 4. TIC-TaC CUT&Tag captures activity dependent remodeling of histone modifications in excitatory neurons. A)** Volcano plot shows excitatory neuron differentially expressed genes (FC > 1.5, Bonferroni-corrected P<0.05) identified by pseudobulk analysis in seizure compared to home cage control nuclei. **B)** Venn diagrams show overlap between downregulated genes identified by MAST (top, blue) and upregulated genes (bottom, red). **C)** Feature plots showing normalized expression in excitatory neurons for representative upregulated gene *H3f3b* (top) and downregulated gene *Htr1f* (bottom). **D)** Violin plots show CUT&Tag normalized reads for H3K9ac binned by downregulated (left, n=158), and upregulated genes (right, n=81). Statistics are two-sided Wilcoxon ranked-sum tests; ****p<0.0001. **E)** Violin plots show CUT&Tag normalized reads for H3K4me3 binned by downregulated (left, n=158), and upregulated genes (right, n=81). Statistics are two-sided Wilcoxon ranked-sum tests; ****p<0.0001. **F)** Gene set enrichment analysis (GSEA) for H3K27ac, H3K9ac, H3K4me3, and H3K27me3. Upregulated and downregulated gene sets were tested for enrichment against histone modification ranked lists ordered by log2 fold change (seizure/home cage). *padj<0.05, ***padj<0.001. **G)** Heatmap shows excitatory neuron downregulated genes (n=158) in seizure compared to home cage control nuclei. Log2 fold changes for gene expression, H3K27ac, and H3K27me3 are shown. Bars to right of heatmap indicate direction of histone modification change (1.2-fold change, teal: H3K27ac decrease, purple: H3K27me3 increase). Pie chart shows percentages of downregulated genes that have decreased TSS H3K27ac only (green), increased TSS H3K27me3 (purple), decreased TSS H3K27ac and increased TSS H3K27me3 (orange), or no change in H3K27ac or H3K27me3 (grey). **H)** Heatmap shows excitatory neuron non DEGs (n=1,016) in seizure compared to home cage control nuclei. Log2 fold changes for gene expression, H3K27ac, and H3K27me3 are shown. Bars to the right of heatmap and pie chart indicate direction of H3K27ac and H3K27me3 changes. **I)** Representative CUT&Tag gene tracks show downregulated (left: *Adcy8, Arhgap15*) and upregulated (right: *Dkk3, Wipf3*) genes. H3K27ac, H3K9ac, H3K27me3, and H3K4me3 are shown for each gene. Data are merged across biological replicates (n=2 home cage, n=3 seizure).

**Supplementary Figure 5. TIC-TaC CUT&Tag captures activity dependent remodeling of histone variants in excitatory neurons. A)** Heatmap shows excitatory neuron non DEGs (n=1,016) in seizure compared to home cage control nuclei. Log2 fold changes for gene expression, H2BE, and H2A.Z are shown. Bars to the right of heatmap and indicate direction of H2BE and H2A.Z changes. **B)** Pie chart shows categories of H2BE and H2A.Z changes among non-DEGs (n=1,016). **C)** Genome browser tracks show previously reported fear conditioning induced changes in H2A.Z. Seizure induced changes at *Egr1* (decreased, left) and *Ppp3ca* (increased, right). Data are merged across biological replicates (n=3 home cage, n=3 seizure). **D)** Heatmap shows excitatory neuron downregulated genes (n=158) in seizure compared to home cage control nuclei. Log2 fold changes for gene expression, H2BE, H2A.Z, H3K27ac, and H3K27me3 are shown. Bars to right of heatmap indicate direction of histone modification or histone variant change (1.2-fold change, green: H2BE decrease, maroon: H2A.Z increase, teal: H3K27ac decrease, purple: H3K27me3 increase). **E)** Heatmap shows excitatory neuron upregulated genes (n=81) in seizure compared to home cage control nuclei. Log2 fold changes for gene expression, H2BE, H2A.Z, H3K27ac, and H3K27me3 are shown. Bars to right of heatmap indicate direction of histone modification or histone variant change (1.2-fold change, green: H2BE increase, maroon: H2A.Z decrease, teal: H3K27ac increase, purple: H3K27me3 decrease).

