## Supplementary Figures for "TEV Induced Cleavage of Nuclei Tagged in Specific Cell Types (TIC-TaC) reveals histone profiles of activated neurons"

### Supplementary Figure 1

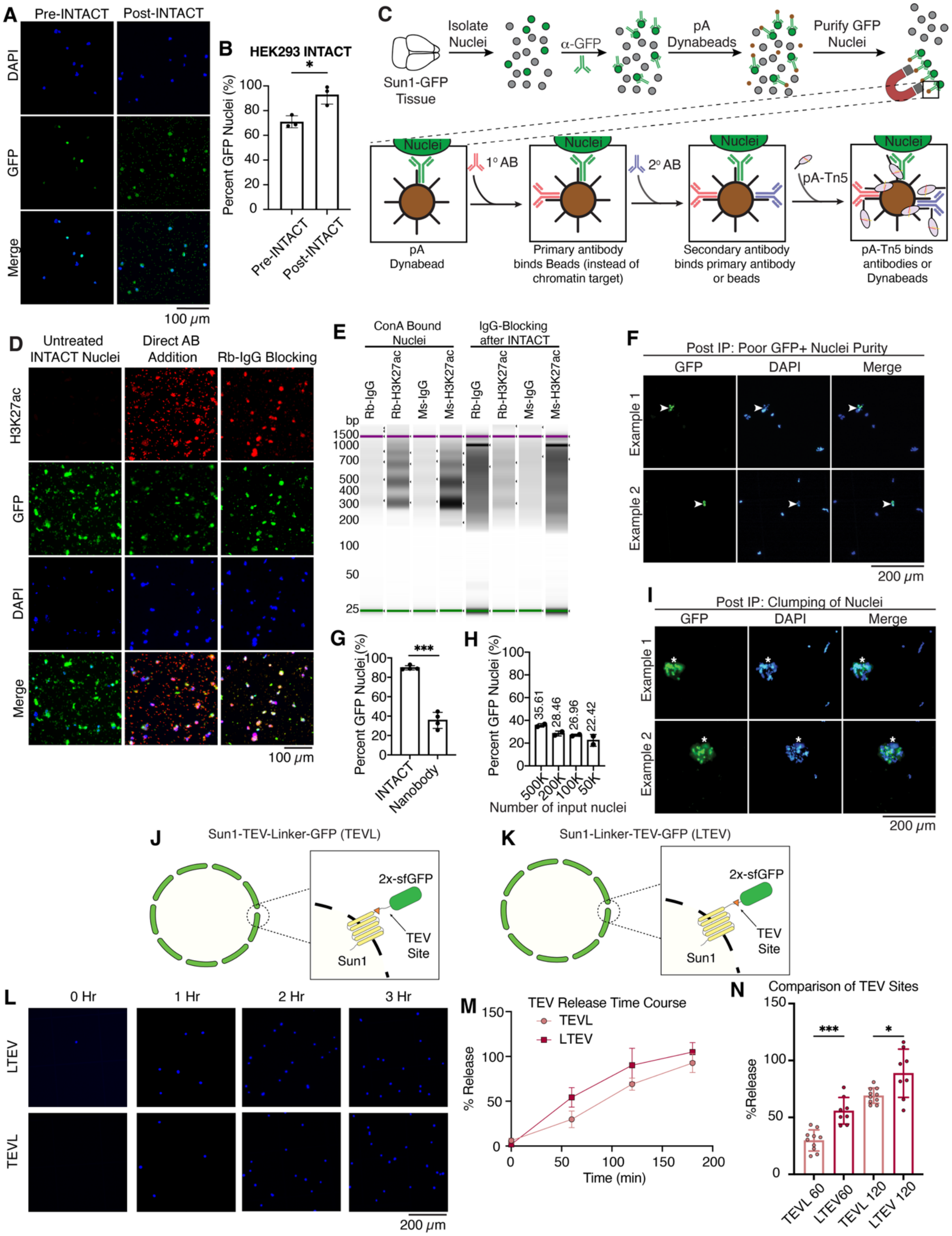

**Supplementary Figure 1. TIC-TaC principle and validation. A)** Fluorescence imaging of pre-INTACT and post-INTACT purified nuclei from HEK293 cells transfected with pCDNA.3_Sun1-GFP, GFP (green) and DAPI (blue). **B)** Quantification of pre-INTACT and post-INTACT purified nuclei from HEK293 cells transfected with pCDNA.3_Sun1-GFP demonstrating enrichment for GPF positive nuclei. (*, $p < 0.05$). **C)** Schematic depicting challenges of combining INTACT with CUT&Tag. Nuclei bound pA-Dynabeads will result in antibodies binding to pA coating beads rather than intended epitopes. pA-Tn5 will then bind to antibodies on the Dynabeads instead of to any antibodies that reach the intended epitopes. **D)** Immunofluorescence of nuclei bound to pA-dynabeads that are not treated with additional antibodies, that are directly treated with an H3K27ac antibody, and that are blocked with excess Rb-IgG prior to the addition of an H3K27ac antibody. H3K27ac visualized by addition of a AlexaFluor-555 (red), with GFP (green), and DAPI (blue). **E)** Tapestation of CUT&Tag performed with an IgG blocking step in bead bound HEK293 GFP positive nuclei. As a positive control, isolated nuclei prior to INTACT were bound to ConA beads. Prior to CUT&Tag on INTACT purified nuclei, Dynabeads were blocked with either excess Rb-IgG or Ms-IgG. For CUT&Tag using Rb-IgG or Rb-H3K27ac, beads were blocked with excess Ms-IgG. For CUT&Tag using Ms-IgG or Ms-H3K27ac, beads were blocked with excess Rb-IgG. **F)** Fluorescence imaging of pre-INTACT and post-INTACT purified nuclei from TIC-TaC$^{+/-}$; Nex-Cre$^{+/-}$ mice using GFP-nanobody beads demonstrating poor purity. **G,H)** Quantification of fluorescence imaging of pre-INTACT and post-INTACT purified nuclei from TIC-TaC$^{+/-}$; Nex-Cre$^{+/-}$ mice using GFP-nanobody beads demonstrating poor purity compared to INTACT (G, ***, $p<0.001$) and that purity is not improved based on the number of GFP positive nuclei in solution prior to purification (H). **I)** Fluorescence imaging of pre-INTACT and post-INTACT purified nuclei from TIC-TaC$^{+/-}$; Nex-Cre$^{+/-}$ mice using GFP-nanobody beads demonstrating clumping of nuclei. **J,K)** Schematic of two different designs for the TIC-TaC system. The TEV site was placed before the linker (J) or after the linker (K) that separates Sun1 from GFP. **L)** Fluorescence imaging of TEV cleavage time course comparing LTEV and TEVL constructs. Time course was conducted at 4C over the course of 3 hours. A minus TEV sample was used as the 0-hour time point. **M)** Quantification of the TEV cleavage time course comparing LTEV (dark red) and TEVL (light red). Near quantitative release was achieved within two hours and three hours respectively. **N)** Quantification of TEV cleavage time course directly comparing LTEV (dark red) and TEV (light red) at 1 hour and 2 hours which demonstrates that the LTEV configuration results in quicker elution from beads (*, $p<0.05$; ***, $p<0.001$).

### Supplementary Figure 2

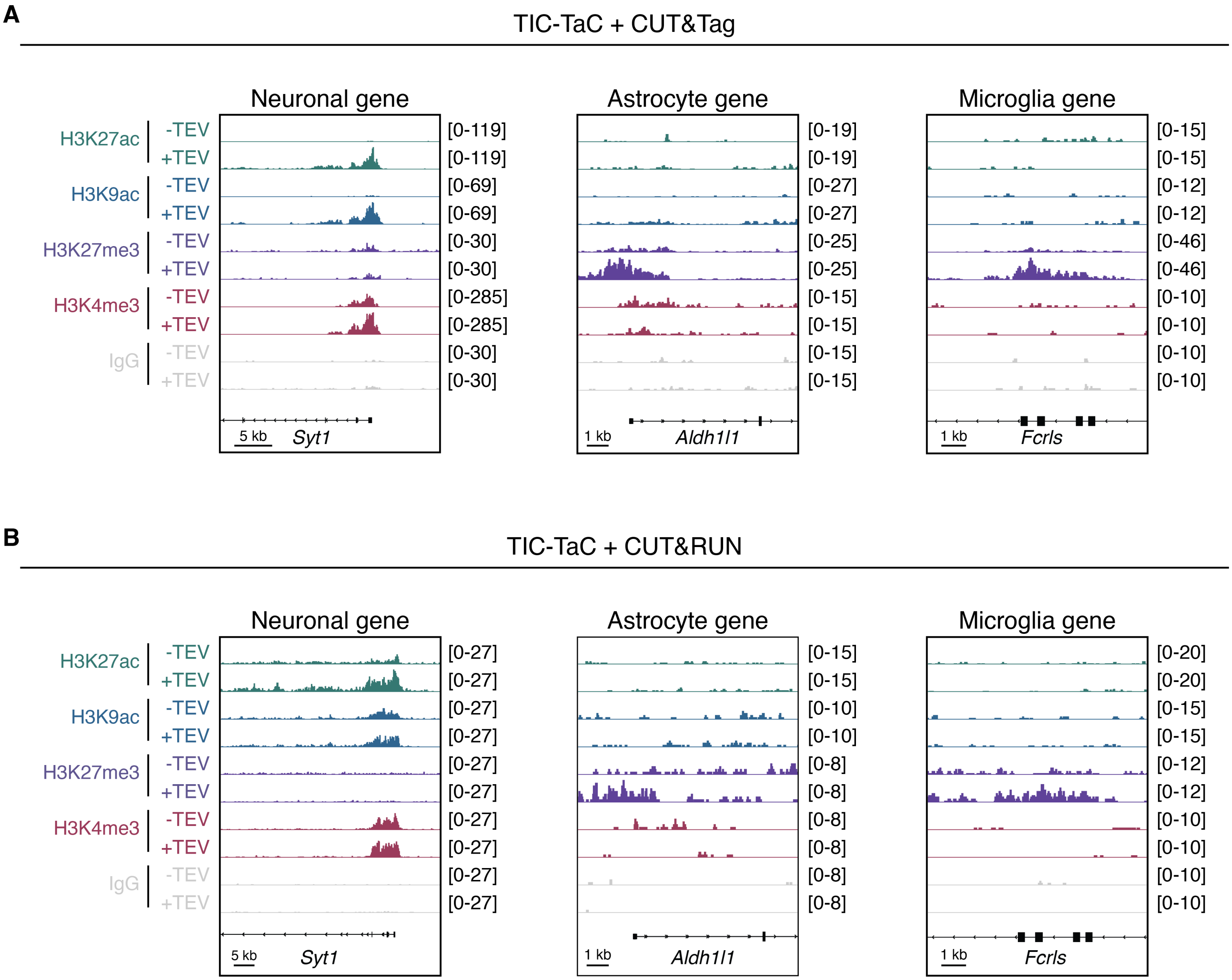


**Supplementary Figure 2. TIC-TaC CUT&Tag/RUN cell-type-specific loci. A)** Representative CUT&Tag gene tracks for a neuronal marker gene (*Syt1*, left) and non-neuronal marker genes (astrocyte: *Aldh1l1*, middle; microglia: *Fcrls*, right). Tracks show n=3 merged biological replicates. **B)** Representative CUT&RUN gene tracks for a neuronal marker gene (*Syt1*, left) and non-neuronal marker genes (astrocyte: *Aldh1l1*, middle; microglia: *Fcrls*, right). Tracks show n=3 merged biological replicates.

### Supplementary Figure 3

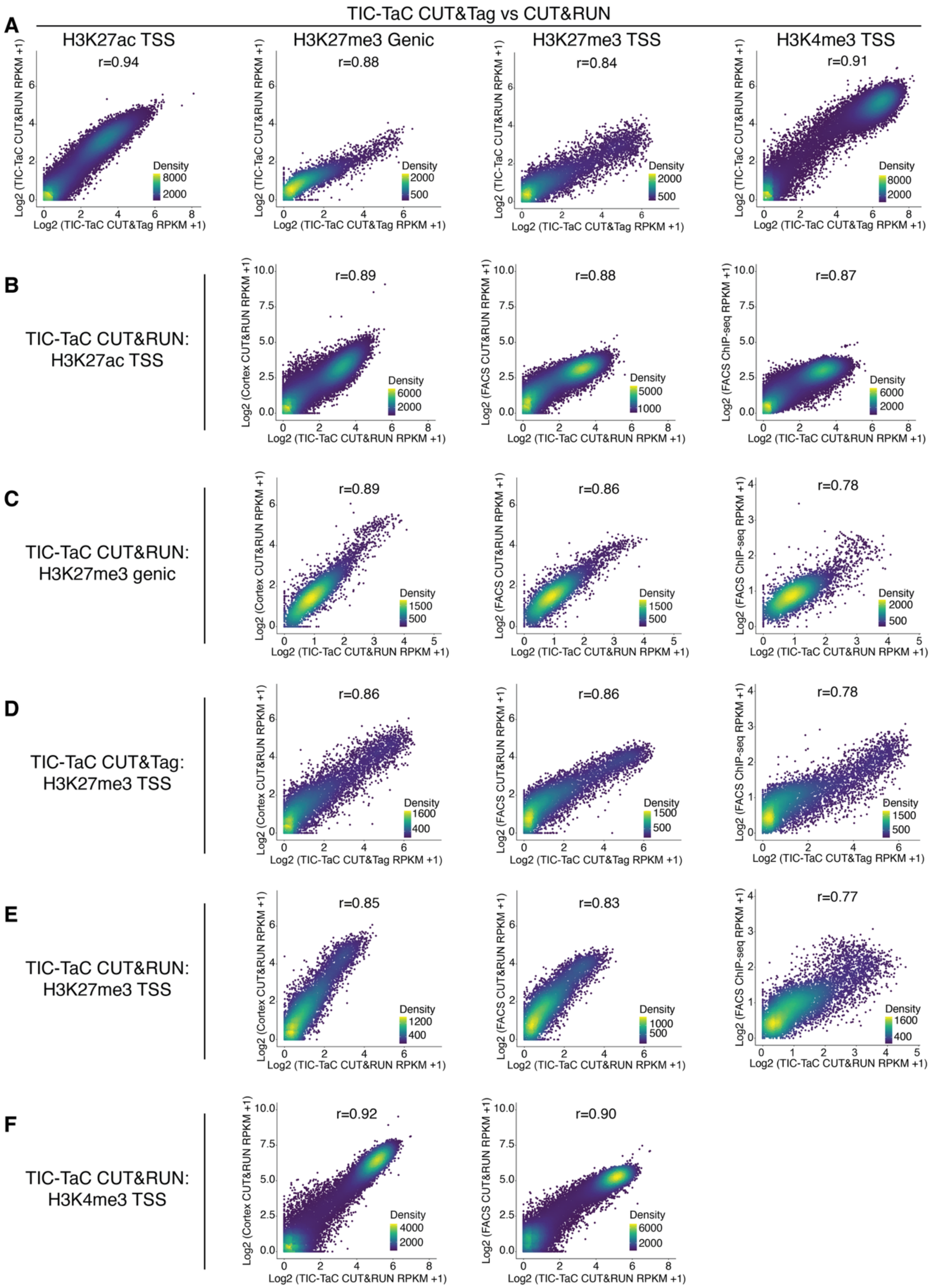

**Supplementary Figure 3. TIC-TaC CUT&Tag/RUN extended benchmarking. A)** Correlation between H3K27ac TSS reads from TIC-TAC CUT&Tag versus TIC-TAC CUT&RUN (Spearman r=0.94, n=29,679), H3K27me3 gene body reads (Spearman r=0.88, n=6,650), H3K27me3 TSS reads (Spearman r=0.84, n=7,084), and H3K4me3 TSS reads (Spearman r=0.91, n=29,679). **B)** Correlation between H3K27ac TSS reads from TIC-TAC CUT&RUN versus bulk cortex CUT&RUN (Spearman r=0.89, n=29,679), versus FACS excitatory nuclei CUT&RUN (Spearman r=0.88, n=29,679), and versus FACS neuronal nuclei ChIP-seq (Spearman r=0.87, n=29,679). **C)** Correlation between H3K27me3 gene body reads from TIC-TAC CUT&RUN versus bulk cortex CUT&RUN (Spearman r=0.89, n=6,243), versus FACS excitatory nuclei CUT&RUN (Spearman r=0.86, n=6,243), and versus FACS neuronal nuclei ChIP-seq (Spearman r=0.78, n=6,243). **D)** Correlation between H3K27me3 TSS reads from TIC-TAC CUT&Tag versus bulk cortex CUT&RUN (Spearman r=0.86, n=7,084), versus FACS excitatory nuclei CUT&RUN (Spearman r=0.86, n=7,084), and versus FACS neuronal nuclei ChIP-seq (Spearman r=0.78, n=7,084). **E)** Correlation between H3K27me3 TSS reads from TIC-TAC CUT&RUN versus bulk cortex CUT&RUN (Spearman r=0.85, n=7,084), versus FACS excitatory nuclei CUT&RUN (Spearman r=0.83, n=7,084), and versus FACS neuronal nuclei ChIP-seq (Spearman r=0.77, n=7,084). **F)** Correlation between H3K4me3 TSS reads from TIC-TAC CUT&RUN versus bulk cortex CUT&RUN (Spearman r=0.92, n=29,679) and versus FACS excitatory nuclei CUT&RUN (Spearman r=0.90, n=29,679).

### Supplementary Figure 4

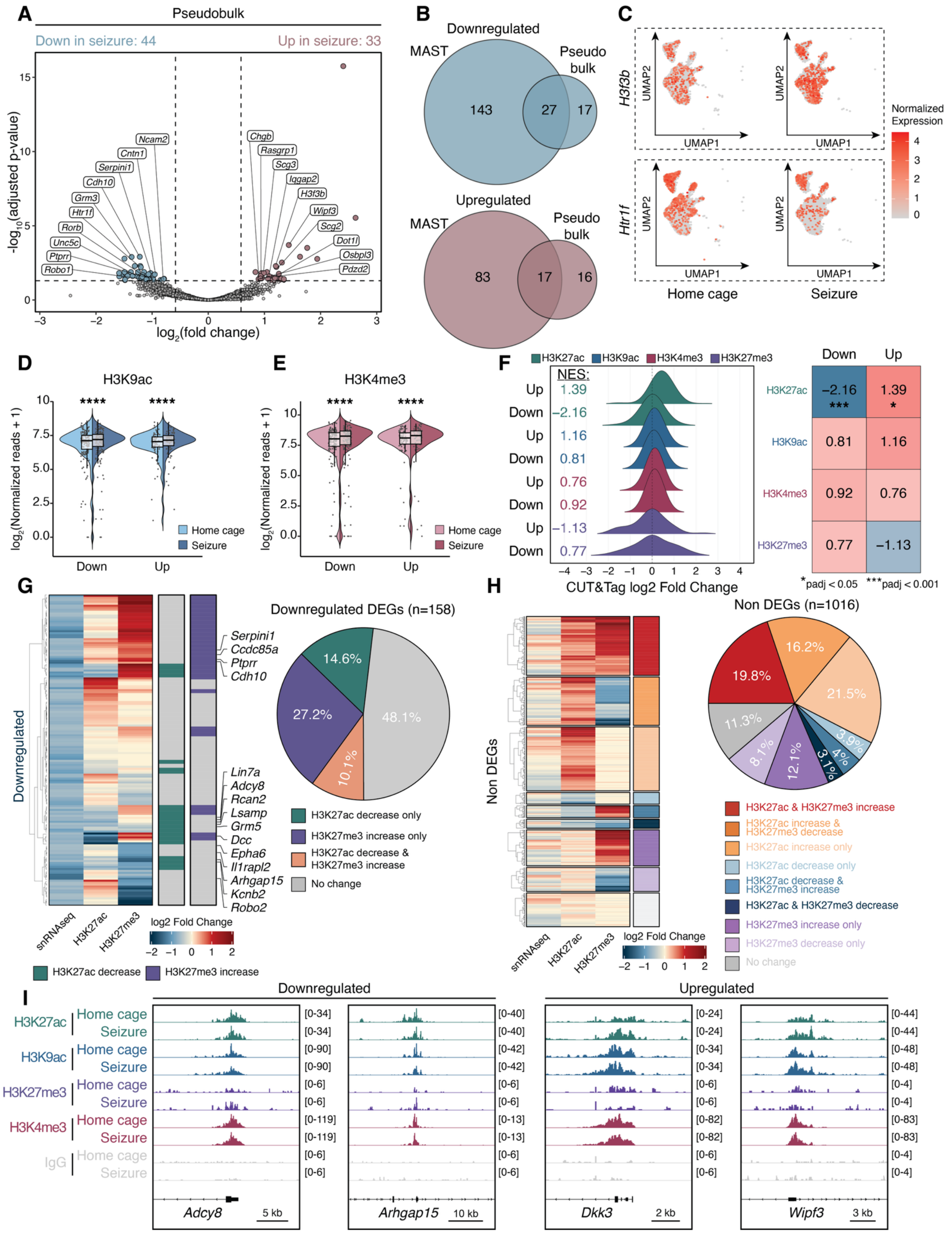

**Supplementary Figure 4. TIC-TaC CUT&Tag captures activity dependent remodeling of histone modifications in excitatory neurons. A)** Volcano plot shows excitatory neuron differentially expressed genes (FC > 1.5, Bonferroni-corrected P<0.05) identified by pseudobulk analysis in seizure compared to home cage control nuclei. **B)** Venn diagrams show overlap between downregulated genes identified by MAST (top, blue) and upregulated genes (bottom, red). **C)** Feature plots showing normalized expression in excitatory neurons for representative upregulated gene *H3f3b* (top) and downregulated gene *Htr1f* (bottom). **D)** Violin plots show CUT&Tag normalized reads for H3K9ac binned by downregulated (left, n=158), and upregulated genes (right, n=81). Statistics are two-sided Wilcoxon ranked-sum tests; ****p<0.0001. **E)** Violin plots show CUT&Tag normalized reads for H3K4me3 binned by downregulated (left, n=158), and upregulated genes (right, n=81). Statistics are two-sided Wilcoxon ranked-sum tests; ****p<0.0001. **F)** Gene set enrichment analysis (GSEA) for H3K27ac, H3K9ac, H3K4me3, and H3K27me3. Upregulated and downregulated gene sets were tested for enrichment against histone modification ranked lists ordered by log2 fold change (seizure/home cage). *padj<0.05, ***padj<0.001. **G)** Heatmap shows excitatory neuron downregulated genes (n=158) in seizure compared to home cage control nuclei. Log2 fold changes for gene expression, H3K27ac, and H3K27me3 are shown. Bars to right of heatmap indicate direction of histone modification change (1.2-fold change, teal: H3K27ac decrease, purple: H3K27me3 increase). Pie chart shows percentages of downregulated genes that have decreased TSS H3K27ac only (green), increased TSS H3K27me3 (purple), decreased TSS H3K27ac and increased TSS H3K27me3 (orange), or no change in H3K27ac or H3K27me3 (grey). **H)** Heatmap shows excitatory neuron non DEGs (n=1,016) in seizure compared to home cage control nuclei. Log2 fold changes for gene expression, H3K27ac, and H3K27me3 are shown. Bars to the right of heatmap and pie chart indicate direction of H3K27ac and H3K27me3 changes. **I)** Representative CUT&Tag gene tracks show downregulated (left: *Adcy8, Arhgap15*) and upregulated (right: *Dkk3, Wipf3*) genes. H3K27ac, H3K9ac, H3K27me3, and H3K4me3 are shown for each gene. Data are merged across biological replicates (n=2 home cage, n=3 seizure).

### Supplementary Figure 5

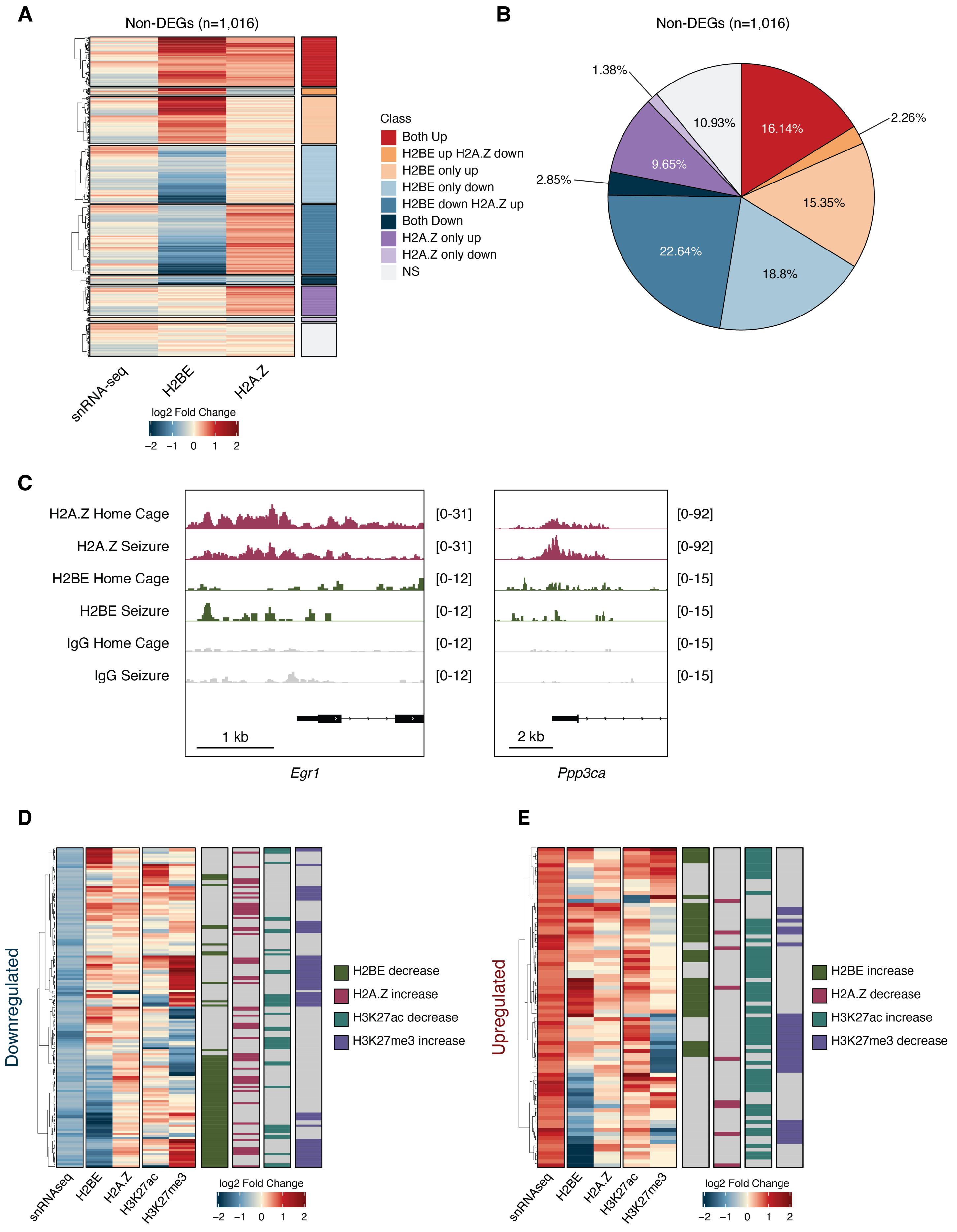

**Supplementary Figure 5. TIC-TaC CUT&Tag captures activity dependent remodeling of histone variants in excitatory neurons. A)** Heatmap shows excitatory neuron non DEGs (n=1,016) in seizure compared to home cage control nuclei. Log2 fold changes for gene expression, H2BE, and H2A.Z are shown. Bars to the right of heatmap and indicate direction of H2BE and H2A.Z changes. **B)** Pie chart shows categories of H2BE and H2A.Z changes among non-DEGs (n=1,016). **C)** Genome browser tracks show previously reported fear conditioning induced changes in H2A.Z. Seizure induced changes at *Egr1* (decreased, left) and *Ppp3ca* (increased, right). Data are merged across biological replicates (n=3 home cage, n=3 seizure). **D)** Heatmap shows excitatory neuron downregulated genes (n=158) in seizure compared to home cage control nuclei. Log2 fold changes for gene expression, H2BE, H2A.Z, H3K27ac, and H3K27me3 are shown. Bars to right of heatmap indicate direction of histone modification or histone variant change (1.2-fold change, green: H2BE decrease, maroon: H2A.Z increase, teal: H3K27ac decrease, purple: H3K27me3 increase). **E)** Heatmap shows excitatory neuron upregulated genes (n=81) in seizure compared to home cage control nuclei. Log2 fold changes for gene expression, H2BE, H2A.Z, H3K27ac, and H3K27me3 are shown. Bars to right of heatmap indicate direction of histone modification or histone variant change (1.2-fold change, green: H2BE increase, maroon: H2A.Z decrease, teal: H3K27ac increase, purple: H3K27me3 decrease).
